# Spatiotemporal control of STING activation and saRNA delivery decouples humoral and cellular immunity

**DOI:** 10.64898/2026.09.11.751026

**Authors:** David J. Peeler, Ziyin Wang, Livia Spiga, Simbarashe Jokonya, Chubicka Thomas, Daniel Reumann, Katsiaryna Shramko, Laia Rigat Nogareda, Paul F. McKay, Dinh Chuong Nguyen, Marco Briones Orta, Patrick S. Stayton, Molly M. Stevens, Robin J. Shattock, John S. Tregoning

## Abstract

Self-amplifying mRNA (saRNA) vaccine formulations link the location and timing of antigen expression to innate immune stimulation by both RNA cargo and carrier. We hypothesized that targeting a polymeric STING agonist prodrug (polySTING) to antigen-presenting cells (APCs) independently of saRNA delivery could tune vaccines responses without limiting saRNA expression. By varying the time and location of STING signaling relative to saRNA delivered by muscle-restricted (polyplex, PP) and lymph node-draining (lipid nanoparticle, LNP) saRNA formulations, we studied how spatiotemporal colocalization of type I interferon (IFN-I) and antigen expression impacts innate and adaptive vaccine responses. Co-administering PP + polySTING decreased antigen-specific adaptive type 1 responses by mismatching early lymph node IFN-I and later arriving antigen; this could be reversed by delaying polySTING delivery timing. Conversely, synchronizing early LNP saRNA delivery and polySTING IFN-I in the lymph node boosted saRNA type 1 T cell responses without impacting humoral immunity, which re-directed antigen-intrinsic immune skewing and enhanced protection against pulmonary *Acinetobacter baumannii* challenge. Collectively, these findings define formulation-specific innate immune responses to saRNA vaccines and establish APC-targeted adjuvants as a strategy to tune saRNA vaccine immunogenicity.

## Introduction

Self-amplifying mRNA (saRNA) enables protective vaccination at low doses, improving both cost effectiveness and overall therapeutic index compared to conventional mRNA vaccines. For example, the first clinically approved saRNA vaccine is dosed at 5 µg, 6-20-fold lower than mRNA vaccines approved to prevent the same infection^1^. Recent clinical studies demonstrated protective seroconversion with just 0.1 µg doses of a rabies saRNA vaccine^2^, effectively increasing the therapeutic index of mRNA vaccines from ∼3 to >100. Tuning innate stimulation to optimize the specificity, potency, and safety of adaptive immune responses remains one of the central objectives in saRNA vaccine design^3^. Besides the ongoing evolution of saRNA construct design^4,5^ and manufacturing^6^, understanding how formulation influences saRNA-induced immune responses is central to the progress of holistic immunoengineering strategies.

Drug delivery vehicles exert direct (via material-intrinsic properties) and indirect (by dictating biodistribution) influence on RNA vaccine immunogenicity and tolerability^7^. Ionizable lipid nanoparticles (LNPs) provide efficient RNA delivery to muscle- and lymph node-resident antigen presenting cells (APCs) while adjuvanting innate responses through pattern recognition receptor binding and/or endosomal damage^8–12^. LNPs driving co-localization of inflammatory signaling (e.g. IFN-I, IL-1β)^13,14^ and antigen expression in lymph node dendritic cells^15,16^ are known to enhance Tfh/B cell interactions in germinal centers. However, while LNP biodistribution to lymph nodes potentiates their effectiveness as RNA vaccines, further biodistribution into systemic compartments contributes to reactogenicity^17^. In contrast, restricting RNA biodistribution to the local injection site with cationic LNPs^18^, lipoplexes^19^, nanoemulsions^20^ or polyplexes (PP)^21–23^ has been associated with lower systemic cytokine and reactogenic responses^24^, but productive immunogenicity varies widely between formulations. For example, we have shown that PP-saRNA formulated with poly(4-amino-1-butanol-co-cystaminebis[acrylamide]) (pABOL) induces 10-fold higher protein expression and 100-fold lower antigen specific IgG titers compared to dose-matched LNPs^25^. The immunological mechanisms underpinning this tradeoff are poorly defined at both the molecular and cellular level, but it is clear that local transfection accesses alternative innate activation and antigen processing pathways compared to formulations that rapidly access the lymph node^13,16,26^. Direct comparisons of innate responses to lymph node-targeted and locally expressed RNA vaccines could thus provide insight into mechanisms to enhance adaptive immunity without reactogenicity.

Type I interferon (IFN-I) is a key factor governing response to RNA vaccination. The magnitude, timing, and location of IFN-I signaling determine whether it enhances or impairs RNA vaccine efficacy. Antiviral IFN-I/IFNAR signaling dampens vaccine responses by restricting saRNA antigen translation in transfected cells^27,28^. Indeed, transient IFNAR silencing in APCs transfected with LNP-saRNA prolongs antigen expression and augments adaptive immunity^29^. IFN-I signaling is, however, essential for dendritic cell activation, T helper 1 (T_H_1) polarization, and cytotoxic T cell proliferation, all of which are critical for durable anti-viral and anti-tumor immunity^30^. The relative timing of IFN-I and antigen presentation further dictates T cell function: simultaneous exposure induces proliferation but IFN-I prior to antigen induces an antiproliferative or proapoptotic state^31^. Collectively, these results suggest that selective activation of IFN-I in lymph node APCs at early timepoints could enhance Type 1 adaptive responses without inhibiting translation or T cell activation. We thus hypothesized that boosting early IFN-I production in lymph node APCs could enhance adaptive immune responses to saRNA vaccines.

Co-formulation of small molecule agonists of the STING (stimulator of interferon genes) pathway with LNP-mRNA has been repeatedly shown to amplify IFN-I signaling, T_H_1 and CD8C T cell responses, and anti-viral/tumor protection^32–40^. Two brief reports have explored unformulated c-di-AMP as an adjuvant for lipoplex and polyplex-delivered saRNA^41,42^, but did not compare to non-adjuvanted saRNA. To define the impact of STING agonism on saRNA expression and immunogenicity, we utilized polySTING, a mannose-targeted polymeric di-ABZI prodrug, to selectively agonize the STING pathway within APCs^43^. In addition to improving potency and safety relative to soluble STING agonists, we hypothesized that polySTING’s hydrophilic chemistry would permit orthogonal co-administration with LNP or PP formulations, thus activating APCs independent of RNA biodistribution. By systematically varying RNA carrier composition and the time and place STING agonism, we sought to define how the location and kinetics of innate stimulation by adjuvanted RNA vaccines determine antigen expression, inflammation, and immunity. Our results outline a generalizable strategy to coordinate saRNA expression and IFN-I signaling to tune adaptive immune responses without compromising safety.

## Results

### PolySTING safely adjuvants polyplex and lipid nanoparticle saRNA formulations through orthogonal lymph node targeting

We reasoned that polySTING would be ideal for co-administration with saRNA formulations because its macromolecular properties should target lymph node APCs without affecting PP or LNP biodistribution (**Fig 1a**). We began by characterizing potential physical interactions between polySTING and pABOL PP^22,24,25^ and C12-200 LNP^44,45^ saRNA formulations, which are in preclinical and clinical development (ISRCTN11364585), respectively. Dynamic light scattering (DLS) showed that polySTING did not induce aggregation when mixed with PP or LNP (**Fig 1b**). Because DLS size distributions are derived from ensemble-averaged measurements and are poorly resolved for heterogeneous populations^46^ (as observed for PP + polySTING), we performed single-particle analysis using fluorescence correlation spectroscopy to confirm that <1% of rhodamine-labelled polySTING (**Fig S1a)** was bound to PP or LNP in solution (**Fig S1b-c**). Based on prior intravenous and subcutaneous polySTING dose-response data comparing APC activation versus systemic and lymph node toxicity^43,47^, we selected 10 µg polySTING (di-ABZI equivalent dose [eq.]; 116 µg net polymer dose) as a safe but robustly activating dose for intramuscular administration. Fluorescent imaging of dissected lymph nodes confirmed that mixing 10 µg polySTING with 1 µg saRNA formulated in PP or LNP before intramuscular injection in mice did not significantly alter polySTING lymph node accumulation compared to polySTING injected alone (**Fig 1c**). Together, these data suggest that polySTING remains physically orthogonal to saRNA nanoparticles when co-injected.

**Figure 1:**
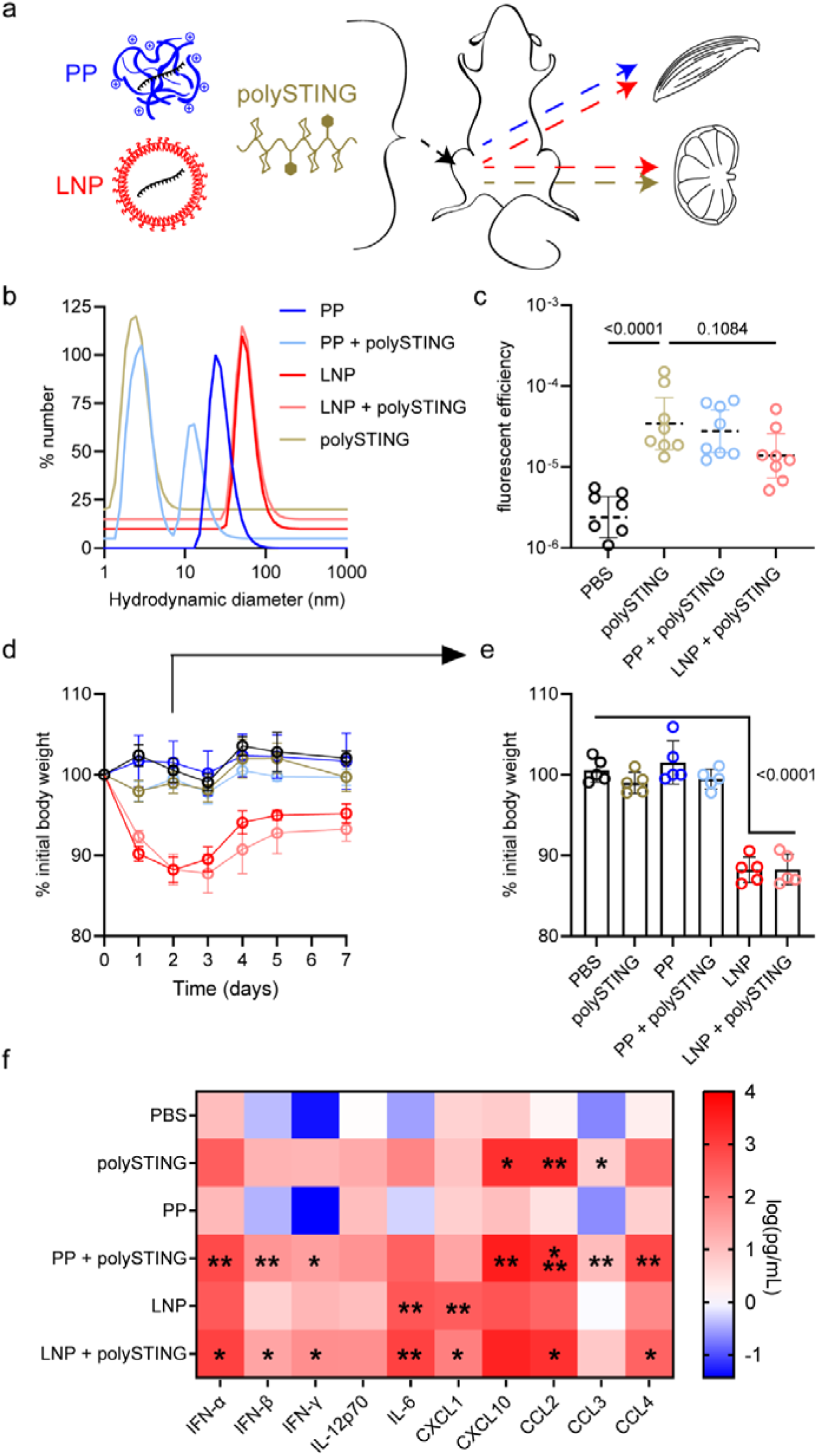
Physicochemical compatibility and complementary inflammatory properties of polySTING, PP and LNP. **(a)** Illustration of hypothesized nanomaterial biodistribution to muscle and lymph nodes. **(b)** Number-weighted hydrodynamic diameter distributions from dynamic light scattering (DLS) of polySTING, saRNA nanoparticles, and their mixtures (*n*=3 measurements; mean shown). **(c)** Fluorescent imaging of polySTING accumulation in inguinal lymph nodes 24 h after intramuscular (IM) injection of 10 µg polySTING (*N*=7-8). After IM 1 µg saRNA ± 10 µg polySTING, **(d)** mean relative body weight over time and **(e)** at day 2 (*N*=5); **(f)** serum cytokine concentration at 6 h (*N*=3-5). Significance by one-way ANOVA with Tukey’s HSD test on log10-transformed values (c) or untransformed values (e), or by Kruskal-Wallis with Dunn’s correction (f). * p<0.05, ** p<0.01, *** p<0.001.

We next assessed how saRNA formulation-intrinsic innate responses were impacted by co-injected polySTING (co-polySTING). Intramuscular administration of polySTING ± PP or LNP saRNA formulations was well tolerated, with no significant polySTING-induced weight loss observed (**Fig 1d**). As previously reported^24^, LNP-but not PP-saRNA induced significant acute weight loss (**Fig 1e**) and systemic inflammatory cytokine levels (**Fig 1f**; **Fig S2a**) including IL-6, which is associated with both reactogenicity and productive humoral responses^7,17^. Correlation analysis showed that all polySTING groups exhibited similar cytokine expression regardless of saRNA formulation (Pearson r^2^ > 0.93, *p* < 1e-4; **Fig S2b**), which consisted of primarily type I and II interferons (IFN-α, IFN-β, IFN-γ) and chemokines (CXCL10, CCL2, CCL3, CCL4). Of note, CXCL1 (a neutrophil chemoattractant) was the only LNP-induced cytokine significantly lowered by co-polySTING yet it remained significantly higher than naïve levels (**Fig S2a**). LNP (and not PP) cytokine expression patterns correlated significantly with their polySTING-adjuvanted counterparts (Pearson r^2^ > 0.69, *p* = 0.026; **Fig S2b**). Elevated cytokine levels largely subsided by 48 h (**Fig S3**). These results show that co-polySTING overlays LNP-like cytokine profiles on that of PP-saRNA without inducing LNP-associated weight loss.

### PP and LNP innate cell recruitment and antigen uptake are differentially modulated by polySTING

We next sought to characterize the acute biodistribution of saRNA transfection and polySTING among the key innate immune cell populations within the injected muscle and draining lymph nodes. Mice received 10 µg (di-ABZI eq.) rhodamine-labelled polySTING ± 10 µg mGreenLantern (mGL) saRNA in PP or LNP in both legs (**Fig 2a**). We selected a higher 10 µg saRNA dose to be consistent with prior biodistribution^20,42,44^ and innate recruitment^48^ studies, where increased RNA input helps overcome the lower sensitivity of fluorescent protein and innate cell counting readouts relative to luminescent reporters and adaptive immunogenicity endpoints. Single cells were dissociated from quadriceps and inguinal lymph nodes and analyzed by flow cytometry 24 h after immunization. PP induced significant innate recruitment to muscle while LNP induced significant innate recruitment to the lymph node (**Fig 2b-c**; gating strategy in **Fig S4**). In muscle, PP primarily recruited neutrophils, macrophages, and NK cells but small numbers of type II conventional dendritic cells (cDC2) were also observed. PP transfection decreased viability among muscle CD3^-^ cells including neutrophils and macrophages, but the addition of polySTING had no additional effect (**Fig S5a**). Co-administration of PP and polySTING decreased NK cell and cDC2 counts in muscle while concomitantly increasing lymph node total innate recruitment by boosting neutrophil, macrophage and type I conventional dendritic cells (cDC1) counts. PolySTING co-administration with PP significantly boosted innate recruitment of every innate cell type studied into lymph nodes to a similar or greater extent than LNP-saRNA only. Co-administration of polySTING with LNP decreased lymph node NK cell, cDC1, and cDC2 counts compared to LNP alone, accompanied by a slight but significant decrease in viability among neutrophils and NK cells analyzed (**Fig S5b**). Interestingly, although polySTING decreased lymph node T cell viability compared to PBS, co-administration with LNP but not PP limited this effect (*p* < 0.0001). These results demonstrate that polySTING re-routes PP innate cell recruitment from muscle to lymph node and antagonizes LNP innate cell recruitment to lymph nodes.

**Figure 2:**
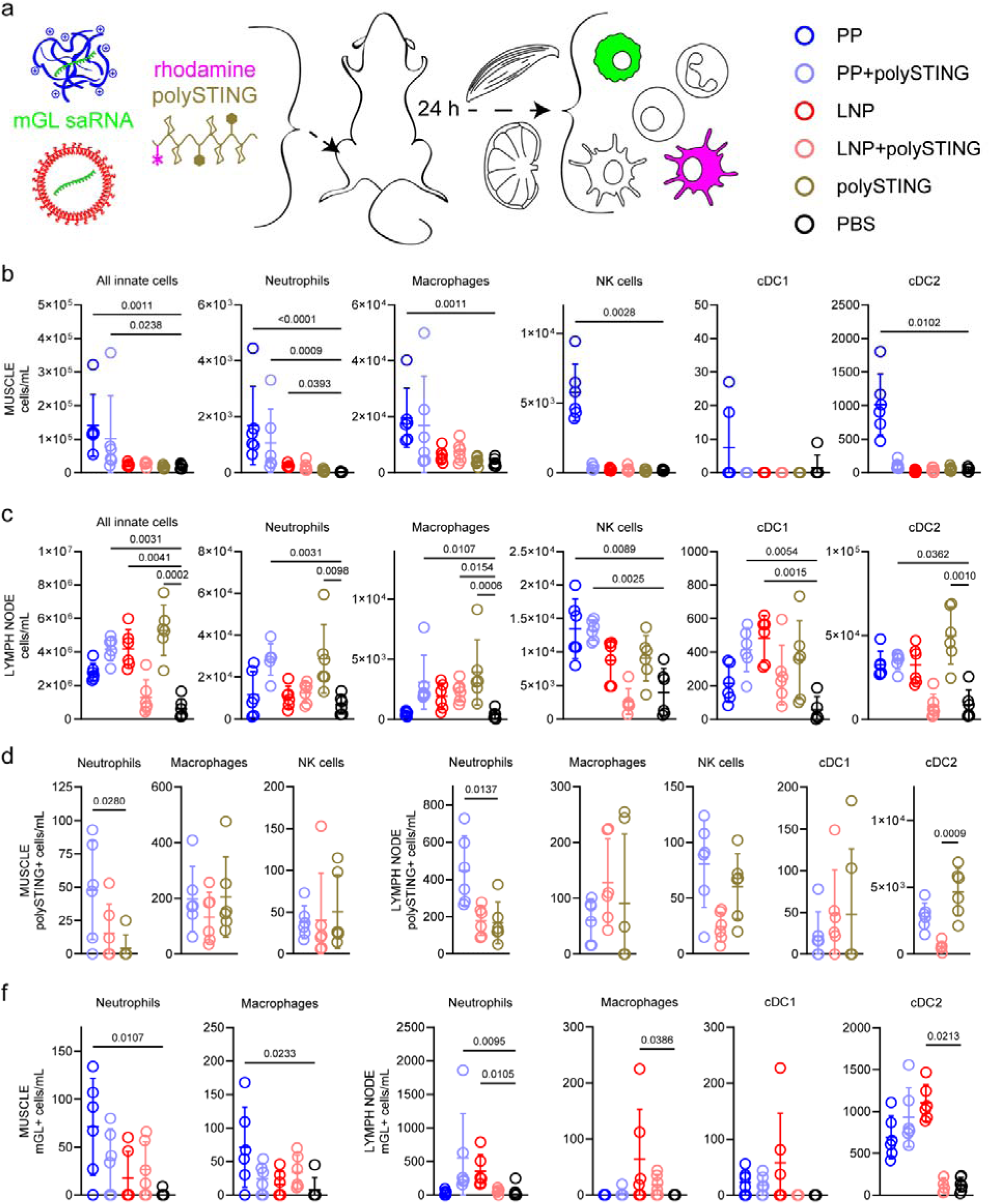
Flow cytometry quantification of acute innate cell recruitment, polySTING uptake, and antigen uptake. **(a)** Mouse quadriceps muscle and inguinal lymph nodes (*N* = 6) were analyzed by flow cytometry 24 h after IM 10 µg mGL saRNA ± 10 µg (di-ABZI eq.) rhodamine-labelled polySTING for: overall cell counts in **(b)** muscles and **(c)** lymph nodes; polySTING+ cell counts in **(d)** muscles and **(e)** lymph nodes; and mGL+ cell counts in **(f)** muscles and **(g)** lymph nodes. Significance by Kruskal-Wallis with Dunn’s correction.

Among the innate cell types analyzed (gating strategy shown in **Fig S6**), we observed rhodamine-polySTING^+^ neutrophils, macrophages, and NK cells in muscle (**Fig 2d**; **Fig S7a**) and rhodamine-polySTING^+^ neutrophils, macrophages, NK cells, and DCs in lymph node (**Fig 2e**; **Fig S7b**). Following PP delivery, macrophages were the most abundant polySTING^+^ cell type in the muscle, whereas cDC2 predominated in the lymph node. While co-injection of PP or LNP with polySTING changed the absolute number of polySTING^+^ cells, it did not change the relative polySTING^+^ frequency within each subtype save for a significant decrease among muscle macrophages and NK cells (**Fig S7b**). Thus, the identity and number of polySTING^+^ cells was largely driven by saRNA formulation-induced innate cell recruitment.

We quantified mGL^+^ innate cells, which include both transfected cells and those that phagocytosed mGL. We found that following PP immunization mGL^+^ macrophages and neutrophils were predominantly observed within the muscle (**Fig 2f**). Although mGL^+^ neutrophils and macrophages were observed in muscle following LNP immunization (**Fig S7c**), most mGL+ cells were detected in the lymph node (**Fig 2g**; **Fig S7d)**, predominantly neutrophils, macrophages, and dendritic cell subsets, with the highest frequencies in cDC1 (∼4%) and cDC2 (∼10%). Co-administration of PP or LNP with polySTING did not affect mGL^+^ frequency within the muscle but decreased the frequency of LNP-driven mGL^+^ cell types in the lymph node.

While dual polySTING^+^/mGL^+^ cells were rare (**Fig S7e-f**), we assessed whether STING agonism directly affected LNP-saRNA translation using RAW264.7-Dual^TM^ reporter macrophages (**Fig S8**). In contrast to mRNA^49^, no one has investigated the impact of STING agonism on saRNA expression. PolySTING increased interferon response factor (IRF) expression in a dose-dependent manner and decreased fLuc saRNA expression by ∼50% at dose ratios >2:1 diABZI:saRNA (w/w). STING-mediated inhibition of saRNA translation, although incomplete, could contribute to the paucity of observed mGL^+^/polySTING^+^ cells.

To sensitively detect the spatial distribution of saRNA expression, we transfected Ai14 mice with 10 µg (di-ABZI eq.) polySTING ± 5 µg Cre saRNA in PP or LNP. We visualized Cre-transfected cells (which become constitutively tdTomato^+^ in the Ai14 line) by performing spinning disk confocal microscopy on tissue-cleared muscles and lymph nodes harvested 24 h after transfection (**Fig 3a**). Although quantification was hindered by autofluorescence, we observed broad PP and LNP transfection throughout the muscle that was unaffected by polySTING co-administration (**Fig 3b**). Transfection appeared to be restricted to interstitial mononuclear cells between myofibers at this early timepoint (**Fig 3c**), consistent with reports for mRNA^26^, which we further confirmed with standard confocal microscopy on cryosectioned tissue (**Fig S9a**). As in our flow cytometry data, we observed markedly greater transfection in lymph nodes for LNP than PP (**Fig 3d**; **Fig S9b**), and total lymph node volume reflected trends in cellular infiltration (PP + polySTING [1.61 mm^3^] ≈ LNP [1.54 mm^3^] >> PP [0.86 mm^3^] ≈ LNP + polySTING [0.85 mm^3^] > PBS [0.65 mm^3^]). These acute data show that polySTING differentially impacts formulation-driven lymph node innate cell recruitment, enhancing PP-driven recruitment but suppressing LNP-driven recruitment, without obviously altering overall transfection biodistribution.

**Figure 3:**
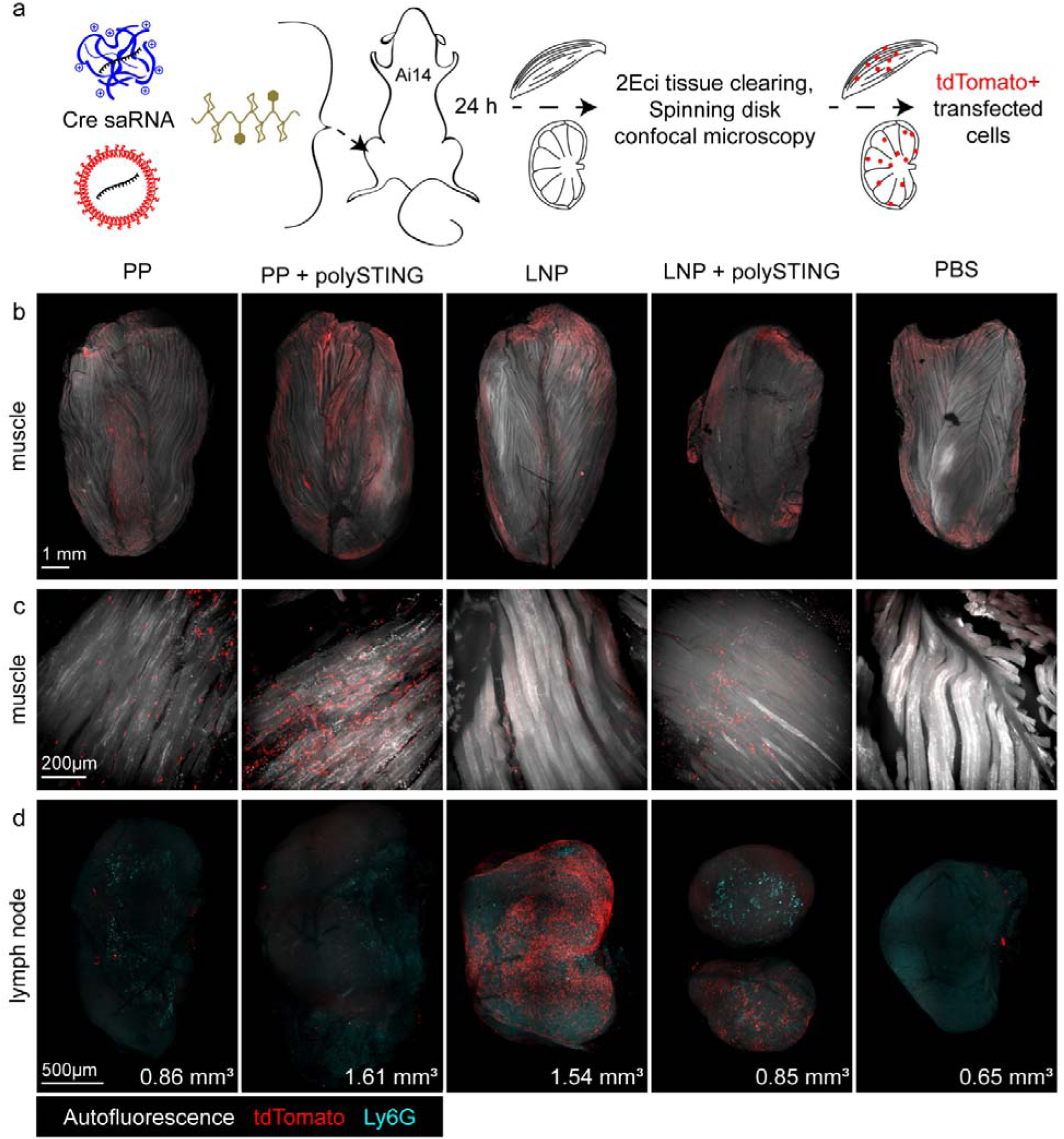
Spinning disk confocal microscopy of transfection in tissue-cleared muscle and lymph node. **(a)** Quadriceps muscle and inguinal lymph nodes (N = 1) were fixed 24 h after IM administration of 5 µg Cre saRNA ± 10 µg (di-ABZI eq.) polySTING and processed for whole-mount imaging using 3D IHC and 2Eci tissue clearing. **(b)** Whole quadriceps imaged using a 4× objective and **(c)** 50 µm z-stack of the same tissue imaged using a 20× objective. **(d)** Z projection of whole inguinal lymph nodes imaged using a 20× objective with total lymph node volume inset at bottom right. Volumes were calculated by trapezoidal integration of cross-sectional area along Z (100 µm steps). Fluorescence channels are indicated in the figure legend.

### saRNA formulation dictates co-polySTING modulation of antigen expression and adaptive immunity

Having seen differential impacts of polySTING on PP and LNP innate immunity, we next asked how polySTING influenced long-term expression and adaptive immunity. Mice were vaccinated bilaterally with 1 µg firefly luciferase (fLuc; right leg) and 1 µg hemagglutinin (HA; left leg) saRNA in PP or LNP ± 10 µg polySTING to enable indirect correlation of expression and immunogenicity (**Fig 4a**). In agreement with recently reported results^37^, *in vivo* imaging of fLuc luminescence revealed that PP and LNP mediated distinct fLuc expression kinetics (**Fig 4b**): PP expression started lower, peaked higher, and concluded earlier. Surprisingly, polySTING significantly increased total PP fLuc expression over time, while no effect was observed on LNP fLuc expression (**Fig 4c**). This increase was driven by prolonged expression in muscle, where ∼100-fold higher expression was observed on day 14 (**Fig 4d**). Interestingly, at a late timepoint (day 26) when PP expression was undetectable, LNP-saRNA transfection was detected in muscle and (putatively) inguinal lymph node, albeit at ∼1% of peak intensity (6.9 ± 2.8 ×10^6^ at day 26 vs. 7.6 ± 3.6 ×10^8^ at day 7; **Fig 4e**). We also assessed immunogenicity using saRNA encoding a membrane-tethered HA previously shown^25^ to protect mice vaccinated with either PP or LNP from lethal influenza challenge. As previously observed, PP induced ∼20-fold lower anti-HA IgG titers compared to LNP (**Fig 4f**); moreover, the addition of polySTING significantly decreased titers induced by PP but not LNP. PP and LNP elicited similar numbers of IFN-γ^+^ splenocytes, but co-polySTING significantly decreased this for PP and increased it for LNP (**Fig 4g**). These results demonstrate that early innate stimulation by polySTING significantly alters long term saRNA expression and immunogenicity in a formulation-dependent manner.

**Figure 4:**
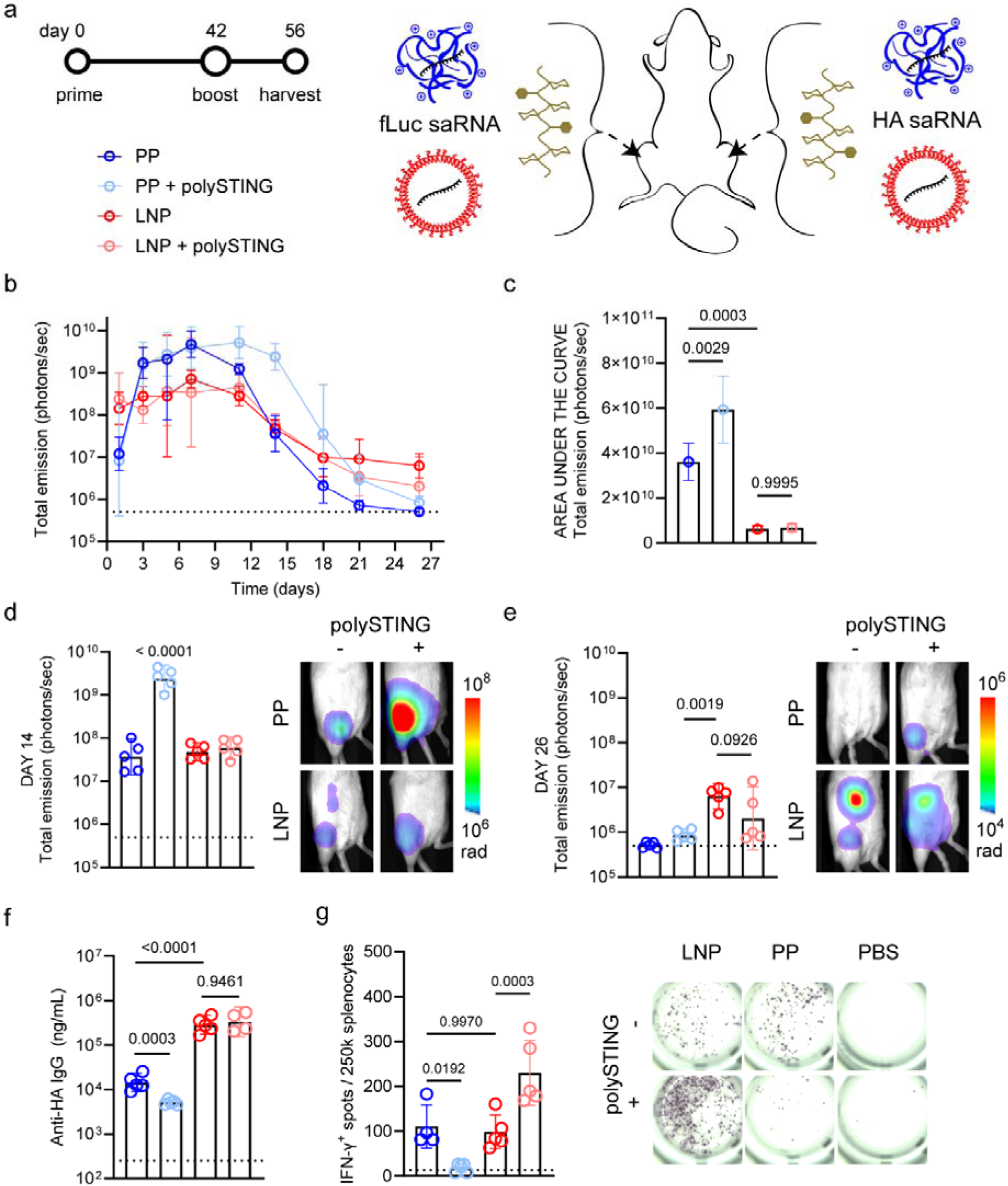
Formulation-dependent effects of co-polySTING on saRNA expression length and adaptive immunogenicity. **(a)** Mice were primed with 1 µg fLuc (right leg) and HA (left leg) saRNA ± 10 µg polySTING, boosted with the same at week 6, and sacrificed for serum and spleen harvest at week 8 (*N* = 5). Luciferase expression **(b)** time course, **(c)** net efficiency (area under the curve), and location at **(d)** day 14 and **(e)** day 26 were analyzed by *in vivo* imaging. Adaptive HA-specific responses were quantified by **(f)** serum IgG ELISA and **(g)** splenocyte IFN-C ELISpot. Dashed line indicates limit of detection; images representative of each group. Significance by ANOVA with Tukey’s HSD test on log10-transformed values (b, d, e, f) or untransformed values (c, g).

### LNP-saRNA + polySTING skews T cell response and reduces disease following bacterial challenge

We next asked whether polySTING could also enhance type I cellular responses against a bacterial antigen encoded in saRNA and non-replicating mRNA constructs. We recently demonstrated that LNP delivery of wildtype (unmodified) mRNA encoding OXA23 better protected mice from lethal intranasal *Acinetobacter baumannii* challenge compared to OXA23 m1ψ-modified mRNA or wildtype saRNA^45^. Thus, we vaccinated mice with 1 µg of LNP-encapsulated wildtype OXA23 mRNA or saRNA either with or without 10 µg polySTING and evaluated disease, bacterial load, antibody titers, and T cell responses in the spleen 1 day after pulmonary challenge with an *A. baumannii* clinical isolate (**Fig 5a**). saRNA + polySTING mediated significant reduction of *A. baumannii* -induced weight loss (**Fig 5b**). Following restimulation with sequential peptide pools spanning the OXA23 sequence, splenocytes from mice vaccinated with saRNA + polySTING generated the greatest IFN-γ^+^ splenocyte responses (**Fig 5c**). Indeed, only saRNA + polySTING resulted in detectable IFN-γ^+^ responses in 5/5 mice vaccinated and the mice protected from weight loss had the highest IFN-γ^+^ spot counts in response to all three peptide pools. While saRNA induced a significant number of IL-17a^+^ spots, the addition of polySTING eliminated this response (**Fig 5d**). Only mRNA alone (7.5 × 10^6^ CFU/mL) and saRNA + polySTING (6.3 × 10^6^ CFU/mL) significantly reduced the geometric mean lung bacterial load compared to unvaccinated mice (1.6 × 10^8^ CFU/mL) (**Fig 5e**). Both polySTING adjuvanted groups failed to significantly decrease spleen infection (**Fig 5f**). Total anti-*A. baumannii* IgG titers were higher for mRNA than saRNA (**Fig 5g**), and while polySTING did not affect total IgG titers for either construct, it decreased mRNA but not saRNA IgG2a (**Fig 5h**). These findings generalize our LNP-HA results to a bacterial antigen and show that STING agonism differentially amplifies saRNA and mRNA immunity.

**Figure 5:**
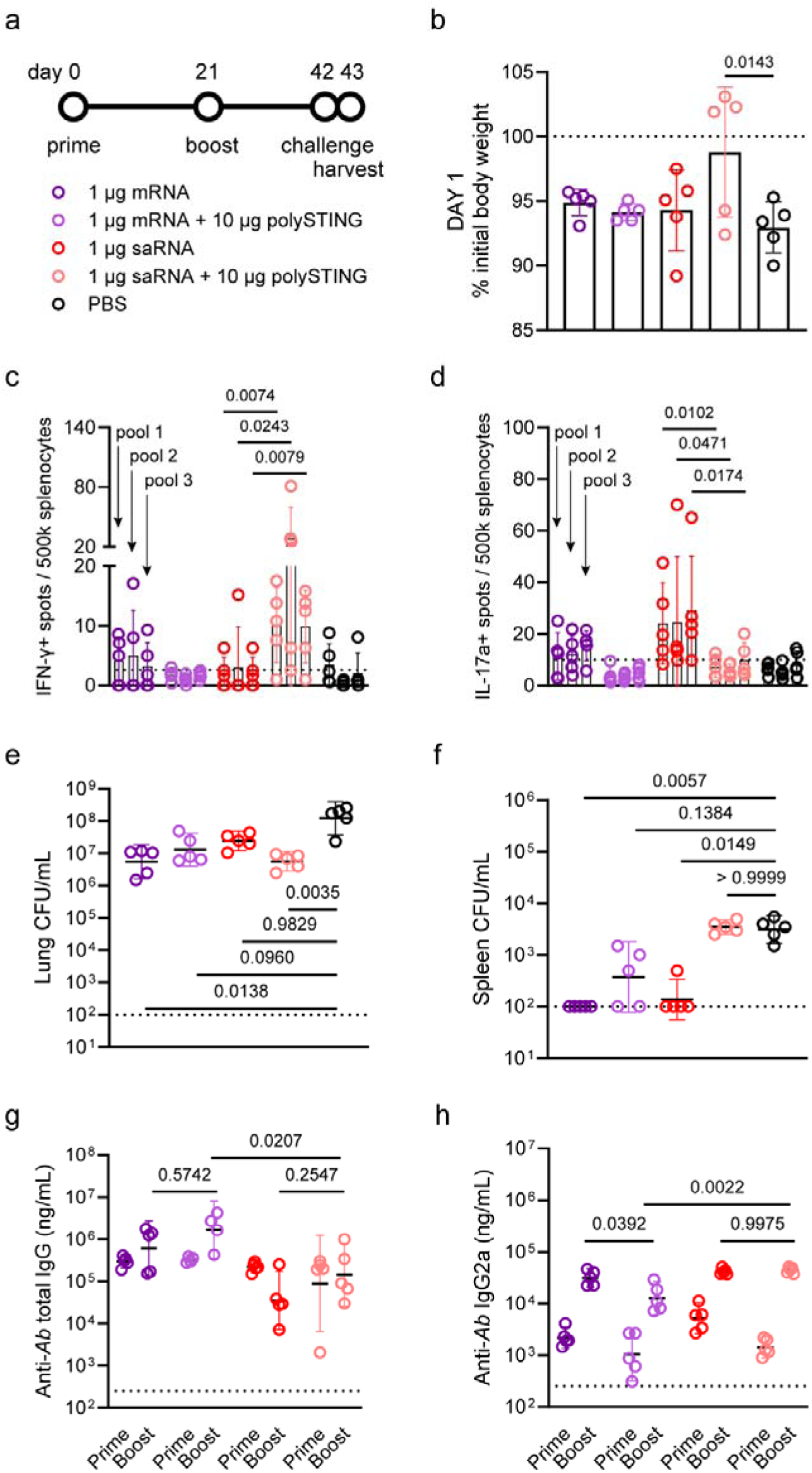
Comparison of polySTING-adjuvanted LNP mRNA and saRNA vaccination in an *Acinetobacter baumannii* challenge model. **(a)** Mice were primed with 1 µg OXA23 mRNA or saRNA ± 10 µg polySTING, boosted in the same way at week 3, intranasally challenged with a clinical isolate of *A. baumannii* at week 6 and sacrificed 1 day later (*N* = 5). **(b)** Infection-induced weight loss was calculated before sacrifice. Cellular adaptive responses were quantified through **(c)** IFN-C and **(d)** IL-17 ELISpot of splenocytes stimulated with OXA23 peptide pools (each dot is the response of one mouse to one pool, where pools are indicated by vertical bars). Bacterial infection was quantified by counting colony-forming units (CFU/mL) in dissociated **(e)** lungs and **(f)** spleens (samples with zero CFU were plotted at the limit of detection). Pathogen-specific humoral responses were quantified through ELISA of serum **(g)** total IgG and **(h)** IgG2a binding to *A. baumannii* outer membrane vesicle lysate. Dashed line indicates limit of detection. Significance by one-way ANOVA with Tukey’s HSD test on untransformed values (b-d) or log10-transformed values (g, h), or by Kruskal-Wallis with Dunn’s correction (e, f).

### Synchronizing lymph node STING agonism and antigen delivery skews humoral immunity

Our observation that synchronizing rapid lymph node delivery of polySTING and LNP-saRNA correlated with increased LNP IFN-γ^+^ splenocyte responses mirrors work where STING agonists were co-formulated with LNP-mRNA^32–40^. Because we observed that PP antigen expression was restricted to muscle, we hypothesized that a temporal mismatch in lymph node innate stimulation (<24 hr) and antigen presentation (>24 hr) was responsible for polySTING-mediated decreases in PP-saRNA adaptive immunity. We thus tested whether decreasing antigen-membrane binding or offsetting polySTING injection could improve PP-saRNA adaptive immunity by synchronizing antigen/adjuvant arrival in the lymph node.

Antigen format (membrane-bound *vs.* secreted) has been shown to affect vaccine IgG induction rate and subtype skew, which is hypothesized to arise from changes in antigen availability, processing, and presentation by APCs^50–53^. To assess whether polySTING-mediated decreases in PP-saRNA adaptive responses were governed by antigen format, we generated a secreted HA saRNA construct (HA_sec_) by removing the transmembrane domain from the parent vector (HA_mem_). We then vaccinated mice with 1 µg HA_mem_ or HA_sec_ PP-saRNA ± 10 µg polySTING and contralateral fLuc PP-saRNA ± polySTING as before (**Fig 6a**). We observed that polySTING-dependent decreases in HA-specific IFN-γ^+^ splenocyte counts were unaffected by antigen format (**Fig 6b**). However, antigen format significantly impacted PP-induced humoral responses, with HA_mem_ eliciting higher total anti-HA IgG (**Fig 6c**; *p* = 0.0402) but substantially lower IgG1 (**Fig 6d**; *p* < 0.0001) and a correspondingly higher IgG2a:IgG1 ratio (**Fig 6f**; *p* < 0.0001) than HA_sec_. When considering both antigen formats, co-formulation with polySTING significantly decreased anti-HA IgG2a (**Fig 6e**; *p* = 0.0174), but not IgG or IgG1. Thus, co-administered polySTING selectively decreases PP T_H_1-associated cellular and humoral responses independent of antigen format.

**Figure 6:**
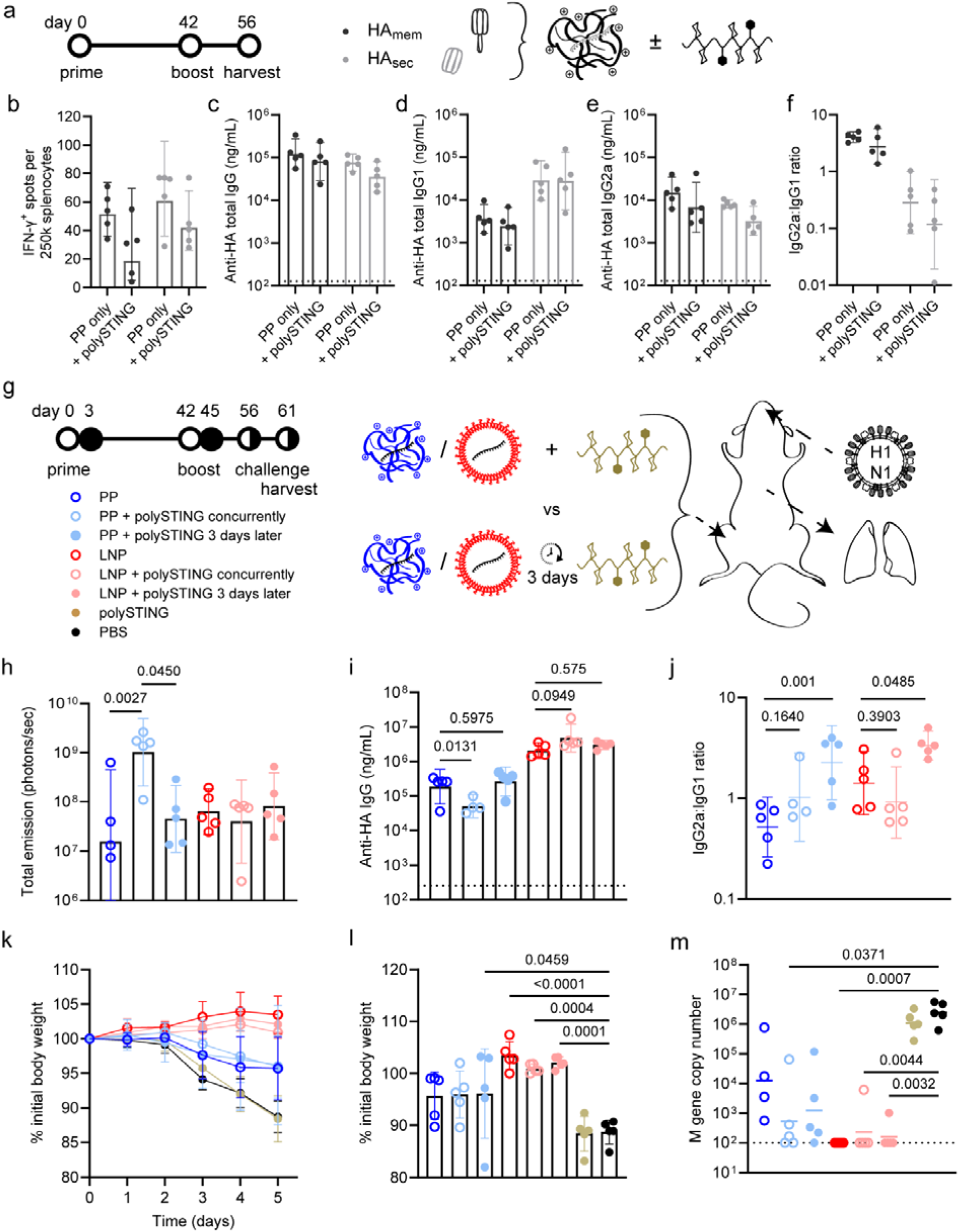
Impact of antigen format and polySTING timing on saRNA immunogenicity and expression. **(a)** Mice were primed with 1 µg fLuc saRNA (right leg) and either membrane bound (HA_mem_) or secreted (HA_sec_) HA saRNA (left leg) ± 10 µg polySTING, boosted with the same at week 6, and sacrificed for serum and spleen harvest at week 8 (*N* = 5). Adaptive HA-specific responses were quantified by **(b)** splenocyte IFN-C ELISpot and serum **(c)** IgG, **(d)** IgG1, and **(e)** IgG2a ELISA; **(f)** the IgG2a:IgG1 ratio was quantified per mouse. **(g)** Mice were primed with 1 µg fLuc (right leg) and HA (left leg) saRNA ± 10 µg polySTING, either in the same injection or given 3 days later. Mice were then boosted in the same way at week 6, intranasally challenged with Cal/09 influenza virus at week 8 and sacrificed for serum harvest 5 days later (*N* = 4-5). **(h)** Luciferase expression on day 13 was analyzed by *in vivo* imaging. Adaptive HA-specific responses were quantified by ELISA of serum **(i)** IgG and **(j)** IgG2a and IgG1. Protection from viral infection was quantified by monitoring **(k)** weight loss over time and **(l)** peak weight loss in addition to **(m)** qPCR of influenza M gene copy number in the lung. Dashed line indicates limit of detection. Significance by one-way ANOVA with Tukey’s HSD test on untransformed values (b, k, l), two-way ANOVA with Tukey’s HSD test on log10-transformed values (c-f), two-way ANOVA with Šídák’s correction on log10-transformed values comparing polySTING timing within each formulation (i,j), or Kruskal-Wallis with Dunn’s correction (m).

We next investigated the impact of delaying lymph node STING agonism relative to transfection. Because transfection-initiated inflammation subsides by 48 hr (**Fig S3**) and antigen expression (**Fig 4b**) continues to increase for both PP and LNP, we hypothesized that delivering polySTING 3 days after transfection would maximally polarize antigen-loaded APCs to induce type 1 adaptive immunity. Using the bilateral fLuc/HA_sec_ experimental design, mice received PP or LNP formulated saRNA without polySTING, with 10 µg co-polySTING, or 10 µg polySTING delivered 3 days later (**Fig 6g**). Concurrent polySTING extended PP fLuc expression (**Fig 6h)**; this was not observed with delayed polySTING. Likewise concurrent (but not delayed) polySTING decreased PP anti-HA IgG titers (**Fig 6i**) as before; polySTING did not impact LNP-delivered saRNA expression or antibody titers. Delayed polySTING delivery induced significant anti-HA IgG isotype switching towards a T_H_1-associated IgG2a bias for both PP and LNP (**Fig 6j**). As observed in previous work^25^, protection from weight loss (**Fig 6k-l**) induced by intranasal viral challenge with Cal/09 H1N1 was driven primarily by the underlying saRNA formulation (PP or LNP). Nevertheless, addition of polySTING to PP reduced geometric mean lung viral titers approximately 11-fold when administered concurrently and 6-fold when administered 3 days later relative to PP alone, and only the concurrent group achieved a significant reduction versus PBS (P = 0.0371), whereas PP alone did not (**Fig 6m**). These data indicate that independent control over antigen and adjuvant pharmacokinetics can tune adaptive immune responses.

### Synchronizing intramuscular STING agonism and antigen expression compromises humoral but not cellular immunity

To investigate whether the location of STING agonism impacts saRNA transfection and immunogenicity, we compared small molecule STING agonists with differing pharmacokinetics to polySTING (**Fig 7a**): di-ABZI (broad bioavailability^54^) or the cyclic dinucleotide ADU-S100 (CDN; lower bioavailability). We also evaluated the effect of co-encapsulation of CDN and saRNA with pABOL (PP-CDN). Building on previous work using the pABOL-like polycation PBAE-447 to encapsulate CDN^55^, we found that pABOL formulation with saRNA:CDN mixtures at 1:1, 2, or 5 w/w ratios resulted in a concomitant increase in PP diameter, indicating co-encapsulation (**Fig S10a-b**).

**Figure 7:**
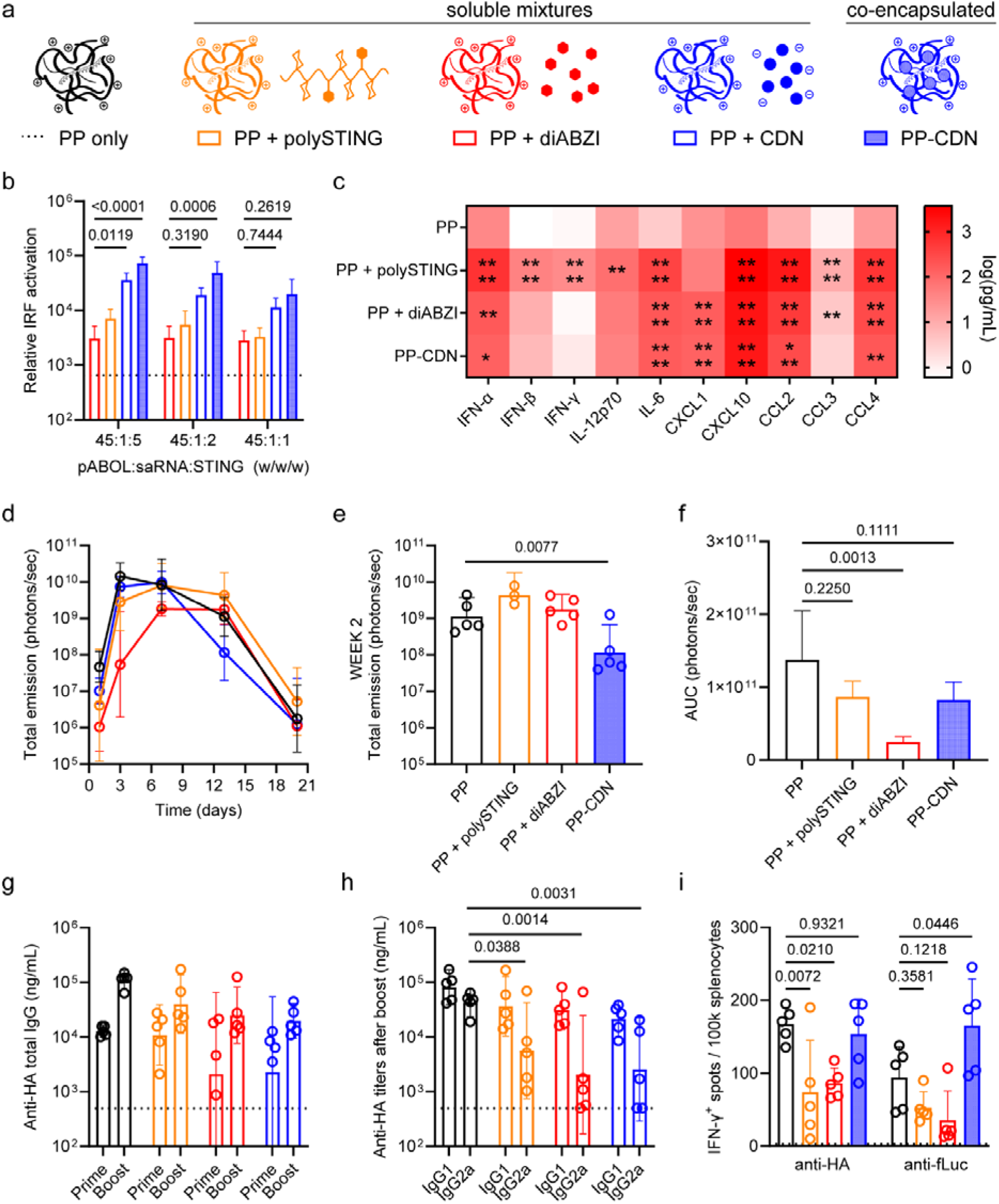
Impact of STING agonist chemistry on PP-saRNA innate activation, expression, and adaptive immune responses. **(a)** Various STING agonists were either mixed with (di-ABZI, polySTING, CDN) or co-encapsulated in (PP-CDN) polyplexes. After 24 h incubation of RAW 264.7 Dual macrophages with 150 ng fLuc saRNA ± various relative STING agonist doses, activation of **(b)** interferon response factor (IRF) was analyzed (*N* = 3). Dashed line shows the mean results of PP only. Mice were injected IM with 5 µg STING agonist mixed with (di-ABZI or polySTING) or co-encapsulated in (PP-CDN) PP delivering 1 µg fLuc (right leg) or HA (left leg) saRNA, boosted with the same at week 6, and sacrificed for serum and spleen harvest at week 8 (*N* = 5). **(c)** Serum cytokine concentration was measured at 6 h (mean shown on log scale for visualization only; statistical comparisons were performed on raw data). The **(d)** time course, **(e)** efficiency at week 2, and **(f)** net efficiency (area under the curve) of luciferase expression were analyzed by *in vivo* imaging. Adaptive responses were quantified by **(g)** HA-specific serum prime/boost IgG, **(h)** boost IgG2a/IgG1 ELISA as well as **(i)** fLuc- and HA-specific splenocyte IFN-C ELISpot. Dashed line indicates limit of detection. Significance by one-way ANOVA with Tukey’s HSD test on untransformed values (b, c, f, i) or log10-transformed values (d, e, g, h). * *p*<0.05, ** *p*<0.01, *** *p*<0.001, **** *p*<0.0001 relative to PP.

When applied to RAW-Dual™ cells *in vitro*, all STING agonists enhanced IRF3 activation in a dose dependent manner compared to PP alone, with PP-CDN (saRNA/CDN co-encapsulation) inducing significantly more activation than admixed PP and CDN, polySTING, or di-ABZI (**Fig 7b**). Only PP-CDN activated NF-κB (**Fig S10c**), and only a slight but insignificant decrease in fLuc saRNA translation was observed at 24 h compared to PP alone (**Fig S10d**).

We thus hypothesized that PP-CDN would colocalize STING activation and transfection in the muscle, analogous to coordinated lymph node STING agonism and transfection by LNP + polySTING. We tested this by performing another HA/fLuc immunization study with 1 µg PP-saRNA ± 5 µg of the STING agonists. While all STING adjuvants significantly increased acute serum levels of IFN-α, IL-6, CXCL10, CCL2, and CCL4 compared to PP alone, only polySTING significantly increased acute IFN-β, IFN-C, and IL-12p70 (**Fig 7c**), demonstrating the enhanced potency afforded by LN targeting. Compared to unadjuvanted PP, fLuc expression was lower for all STING adjuvanted PP at day 1 (**Fig 7d**) but only PP-CDN expression was significantly lower at week 2 (**Fig 7e**) and only PP + di-ABZI resulted in significantly lower net expression (**Fig 7f**). The modest impact of polySTING on transfection extension in this experiment (compared to **Fig 4b-d**) may reflect the lower polySTING dose used. At harvest, we observed that STING agonism decreased anti-HA IgG titers (**Fig 7g**; two way ANOVA, *p* = 0.0369), primarily by reducing IgG2a responses (**Fig 7h**). We further confirmed that STING-mediated decreases in Ig2a titers were not reflective of a skew towards IgE, allergy-linked responses (**Fig S11**). Lymph node-draining STING agonists (di-ABZI, polySTING) decreased anti-HA IFN-γ^+^ splenocyte responses relative to PP alone (**Fig 7i**), an effect that was generally (but not significantly) conserved for anti-fLuc responses. Collectively, these data suggest that concurrent STING agonism generally antagonizes antibody responses to PP-saRNA, but restricting STING agonism to the muscle reduces suppression of T cell immunity.

### Neutrophils are dispensable for polySTING immunomodulation, but IFNAR signaling is not

Having shown that the timing and location of STING agonism could be used to manipulate adaptive humoral and cellular responses to saRNA vaccines, we sought to confirm the key innate cell types and signaling pathways responsible. The recurring prominence of neutrophil signatures in our cytokine, flow cytometry, and microscopy data prompted us to perform a transfection/immunization study in mice with and without systemic neutrophil depletion during prime-boost vaccinations (**Fig S12a**). Comparing mice receiving 1 µg fLuc/HA saRNA delivered by PP, PP + 10 µg polySTING, or LNP, we observed no differences in fLuc expression (**Fig S12b**), anti-HA IgG titers (**Fig S12c**), or HA-specific IFN-γ^+^ splenocyte responses (**Fig S12d**) following anti-Ly6G treatment. Although neutrophil depletion resulted in a slight but insignificant decrease in cellular responses to unadjuvanted PP and LNP, our results indicate that neutrophil depletion has little effect on adaptive responses to saRNA vaccines.

To query the role of IFN-I signaling, we compared innate and adaptive responses in animals receiving prime-boost vaccination with 1 µg HA saRNA in PP or LNP to those receiving PP or LNP with 10 µg polySTING ± systemic IFNAR blockade by IgG clone MAR1-5A3 (**Fig 8a**). For LNP + polySTING, blockade significantly reduced seven of ten serum cytokines, with IFN-γ and CXCL10 approximately fourfold lower, while IFN-β increased by approximately threefold, consistent with a loss of autocrine IFNAR-mediated negative feedback on IFN-β production (**Fig 8b**; **Fig S13**). Both cytokines and chemokines were affected, indicating upstream IFNAR (rather than direct effects of STING signaling) largely governed polySTING-mediated increases. For PP + polySTING, blockade produced a smaller effect confined to cytokines, with IL-6 falling 2.68-fold (*p* = 0.0141). PolySTING enhanced chemokine responses to both LNP and PP delivered RNA, but that induction required IFNAR signaling only after LNP delivery.

**Figure 8:**
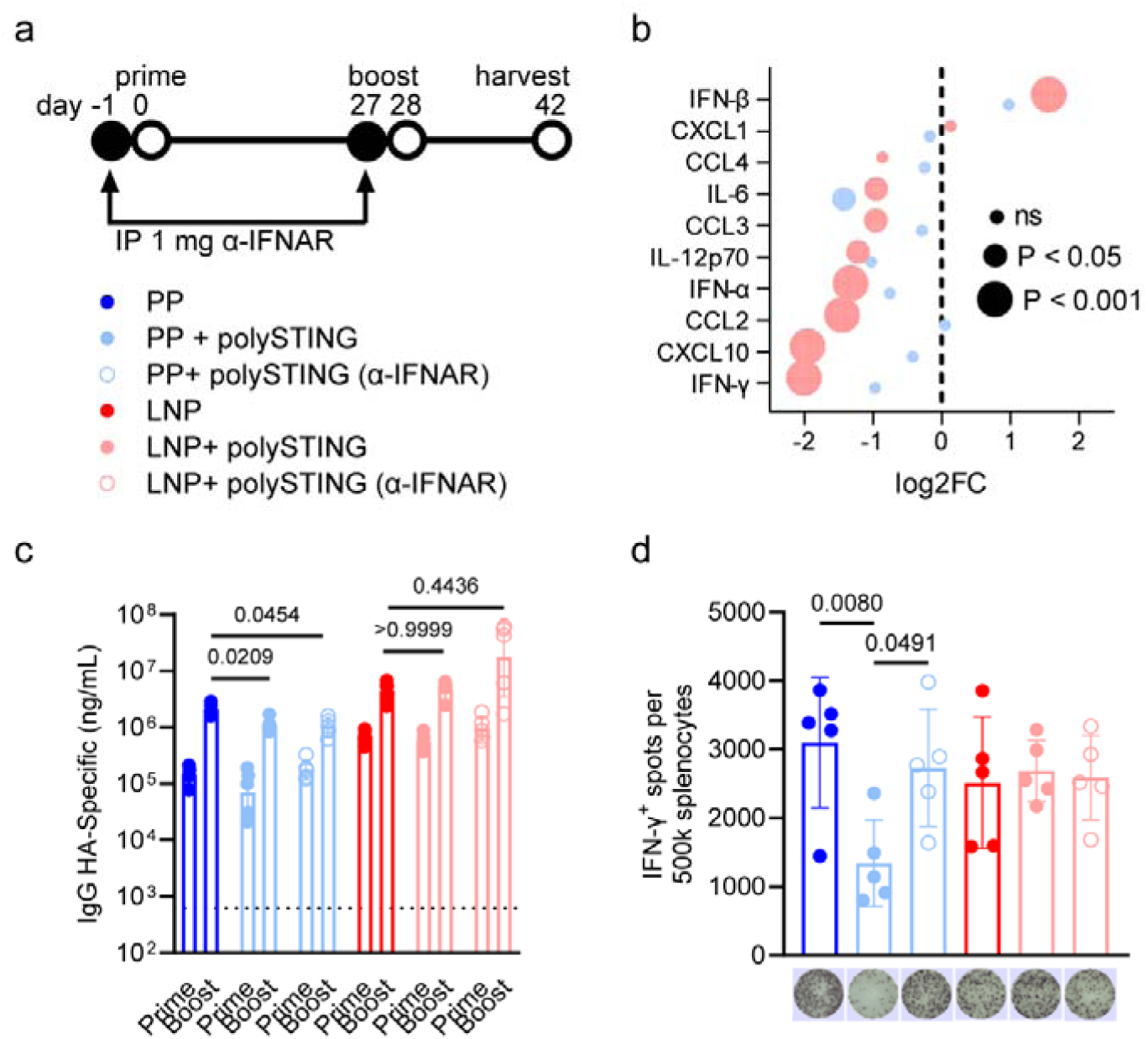
The role of IFNAR signaling in polySTING-modulated transfection and immunogenicity. **(a)** Mice were dosed intraperitoneally (IP) with anti-IFNAR antibodies on the day before and after receiving 1 µg HA saRNA ± 10 µg polySTING according to the prime/boost schedule shown (*N* = 5). **(b)** Serum cytokine levels were quantified at 6 h after priming and plotted as the fold change in mean log2 concentration between the PP (blue) or LNP (pink) + polySTING + αIFNAR group relative to the corresponding PP/LNP + polySTING group. HA-specific adaptive responses were quantified by **(c)** serum IgG ELISA and **(d)** splenocyte IFN-C ELISpot (negative control subtracted from each mouse; pictures of representative wells shown below). Dashed line indicates limit of detection. Significance by two-way ANOVA with Šídák’s correction on log2-transformed values (b) or one-way ANOVA with Dunnett’s correction on log10-transformed (c) or untransformed (d) values.

Consistent with our earlier observations, polySTING reduced PP but not LNP IgG after boost, and this reduction was unchanged by IFNAR blockade, indicating that the humoral deficit does not require IFN-I signaling (**Fig 8c**). PolySTING reduced PP IFN-γC splenocyte frequencies, and IFNAR blockade significantly restored them to levels indistinguishable from PP alone (*p* = 0.9747), indicating a recovery of the T cell response (**Fig 8d**). Neither polySTING nor blockade significantly altered LNP IFN-γC frequencies in this experiment, although we have previously observed polySTING-mediated enhancement of LNP T cell responses under a different prime-boost interval and antigen configuration.

## Discussion

PP and LNP saRNA vaccines each have distinct advantages and limitations that likely arise from how their differing biodistribution and inflammatory properties shape innate and adaptive immune cell engagement. PPs generate high local antigen expression with low reactogenicity but weaker humoral immunity, whereas LNPs induce strong inflammation and robust antibody responses^24^. In contrast to humoral immunity, published comparisons of T cell priming by PP and LNP saRNA formulations are rare^25^ and the intermediate innate mechanisms between local transfection and adaptive immunity are poorly defined^13,26^. Differences in IFN-I induction between these platforms are also likely to shape adaptive cellular immunity since IFN-I is a key regulator of T cell expansion and differentiation^31^. We therefore investigated how an early, STING-mediated IFN-I pulse from APCs affects the location and duration of saRNA expression, innate-cell recruitment, and downstream cellular and humoral immunity across these two delivery platforms. A recent study^47^ demonstrated that separately targeting peptide antigen and polySTING to the lymph node through subcutaneous injection boosted anti-tumor T cell responses. Our data extend this concept into the intramuscular saRNA vaccine space, showing how one can modulate induction of humoral and cellular adaptive immunity depending on the site and timing of antigen expression and STING stimulation.

Given findings that inhibiting IFNAR signaling in saRNA-transfected cells increases and sustains transgene expression^27,29^, it would be intuitive to expect polySTING IFN-I to inhibit saRNA expression. In agreement with recently reported STING-mediated decreases in mRNA expression^49^, we observed that STING agonism decreased *in vitro* saRNA expression by 50% at 24 hr. However, polySTING did not decrease saRNA expression *in vivo*, and to our surprise, significantly extended expression of PP-saRNA. Because we observed a lower level of luciferase-specific IFN-C^+^ splenocytes alongside extended PP-saRNA expression, echoing findings with DNA vaccines^56^, we inferred that polySTING modulates the antigen-specific T cells that eliminate saRNA-expressing cells. Finding that only concurrent (and not delayed) polySTING prolongs PP-saRNA expression implicates early modulation of lymph node T cells with IFN-I as a means to engineer long-term saRNA function.

Spatiotemporally decoupling antigen expression and adjuvant stimulation by coadministering PP + STING agonists enabled us to interrogate how IFN-I timing and location direct adaptive vaccine responses. We demonstrated that lymph node-draining, but not muscle-restricted, STING agonists decreased type I T cell responses to IM PP saRNA vaccines in an IFNAR-dependent manner. While PP-concurrent administration of either local or lymph node-draining STING agonists significantly decreased both IgG and IgG2a titers, delaying polySTING delivery preserved total IgG, increased skew towards IgG2a, and improved local control of influenza lung infection relative to a non-adjuvanted PP HA saRNA vaccine. The T_H_1 skew induced by delayed polySTING reflects other reports that separately administered TLR9 and STING agonists synergize to enhance antigen-specific T_H_1-associated responses and suppress the T_H_2 bias intrinsic to STING agonism alone, an effect attributed to the two adjuvants acting on distinct cell populations rather than to IFN-I^57^. PP-saRNA induced moderate IL-12p70 with little IFN-I (**Fig 1f, 7c**), and IFNAR blockade had only modest effects on PP + polySTING innate responses (**Fig 8b**), making IL-12 a plausible priming signal analogous to TLR-driven MyD88 signaling ^57^. Although we did not measure lymph node cytokines or APC activation at the time of delayed dosing, our flow cytometry and microscopy data clearly define innate cell migration as a key feature of PP immunization. We thus link the polySTING-mediated loss of PP T cell responses to a spatiotemporal disconnect between IFN-I and antigen presentation. Polyplexes transfect predominantly at the injection site, whereas polySTING drives IFN-I production in the draining lymph node, and IFN-I arriving before antigen presentation is known to impair T cell expansion^31^. Productive cellular immunity to saRNA vaccines therefore appears to require coincident antigen and IFN-I signaling, so adjuvants acting on the interferon axis must be matched to the biodistribution of the platform they accompany.

We showed that concurrent polySTING delivery enhanced IFN-γ^+^T_H_1 cellular responses to both viral and bacterial antigens expressed from LNP saRNA without impacting humoral immunity. Synergy between synchronized lymph node delivery of polySTING and C12-200 LNPs (which may stimulate TLR4^11,58^) echoes recent reports of an MPLA/cdGMP dual-adjuvant liposome that boosted IFN-γ^+^ CD4^+^ T cell responses in an IFN-I dependent manner^59^. Using an OXA23 peptide pool, we showed that mRNA and saRNA OXA23 induce T_H_17-biased splenocyte responses similar to previous findings with live attenuated *A. baumannii*^60^ and plasmid-expressed OmpA/Pal^61^ vaccines. Co-administration of polySTING shifted saRNA-induced splenocyte responses from IL-17a^+^ to IFN-γ^+^, reducing disease and lowering lung bacterial burden. These findings agree with prior *A. baumannii* conjugate vaccine studies attributing T_H_1-enhanced protection to IFN-γ mediated macrophage activation, which increased macrophage bactericidal capacity and ultimately reduced T_H_17 polarization^62^. Our data support the emerging role of adjuvanted nucleic-acid vaccines for bacterial pathogens^63^ and indicate that saRNA may offer unique opportunities to harness T_H_1-biased bacterial protection.

Antigen format changed the skew of PP-induced humoral responses, but not cellular responses, nor the impact of polySTING on those responses. Membrane-bound antigens have been reported to induce similar IFN-γ^+^ splenocyte responses^52^ but a higher titer, IgG2a-skewed IgG repertoire^50,52^ compared to secreted formats, whereas secreted antigens induce larger germinal centers with greater numbers of T_FH_ cells^52^. These differences are posited to arise from a combination of passive and active antigen transport mechanisms, as well as the orientation of antigen epitopes when presented to B cells^64^. Our results support a similar skew (IgG2a for HA_mem_, IgG1 for HA_sec_), but do not explain which cells are responsible for antigen trafficking and presentation that give rise to these skews. Given the limited numbers of mGL^+^ cells we observed at the relatively high 10 µg dose used in **Fig 2**, future studies of innate recruitment/antigen trafficking would benefit from advanced methodology such as photoconvertible proteins^13^ or genetically encoded cell-cell mapping techniques^65^ for sensitivity at immunization-matched doses.

We found that polySTING modulated formulation-specific relationships between lymph node innate recruitment and adaptive outputs in a counterintuitive manner. Flow cytometry and microscopy showed that polySTING increased PP-driven but reduced LNP-driven innate cell recruitment to lymph nodes, with correspondingly lower and greater adaptive outputs, respectively. We focused our analyses on the key myeloid (cDCs, macrophages, NK cells) and lymphoid (T cell) cells that express STING ^66^, along with neutrophils, the most abundant polySTING^+^ cell type in PP-transfected muscle (**Fig 2b, 2d**) and the most abundant polySTING^+^ cell type in LNP-transfected lymph node (**Fig 2e**). While neutrophils have been shown to be dispensable for humoral responses to LNP HA mRNA^67^ and repress STING expression under homeostatic condtions^66^, they can function as APCs^68^ and STING signaling in neutrophils can induce strong systemic inflammatory pathologies.^66^ PolySTING decreased LNP-mediated neutrophil chemoattractant expression (CXCL1; **Fig S2-S3**), lymph node mGL^+^ neutrophil frequency (**Fig 2**), and viability (**Fig S5**). We thus hypothesized that polySTING might improve LNP-mediated T cell responses by decreasing neutrophil involvement and thus potentially enabling true APCs to access antigen. Our neutrophil depletion study showed that neutrophils were in fact dispensable for both PP and LNP saRNA vaccine responses. Thus, neutrophils function mainly as a non-functional sink for polySTING and are not essential for saRNA vaccine immunogenicity. Additional mechanistic work is needed to dissect which cells serve as the essential intermediates of STING-enhanced and STING-suppressed T_H_1 responses. For example, the recent finding that IFN-I increases cross-dressing of cDCs^69^ makes it plausible that polySTING directly boosts the efficiency of LNP-initiated antigen presentation by DCs. Conversely, it would be interesting to examine whether polySTING mediates antigen tolerance to PP-expressed antigen through IL-10^+^ T cells and DCs implicated in past work^70^. Despite the established mechanisms of polySTING targeting myeloid cells, our data does not rule out the possibility that other stromal and immune cell types in other organs could contribute to the outcomes observed, especially given their newly discovered roles in antigen presentation^71^.

Several limitations should be considered when interpreting these data given that all *in vivo* studies were performed in female BALB/c mice using mRNA or saRNA composed of wildtype uridine. We^45^ and others^72^ have found a benefit of wildtype relative to m1ψ-modified mRNA in the context of bacterial vaccines, reflecting the different balance of adjuvant activity and antigen expression that the two chemistries establish. Species-specific differences in pattern recognition receptor sensing may merit further comparison of these nucleotide chemistries during clinical testing (e.g. NCT05537038 *vs.* NCT05547464). While BALB/c mice are commonly associated with T_H_2-skewed immunity, we^45^ and others^36^ have shown that mRNA-LNP vaccination can still induce robust T_H_1-associated IgG2a and cellular responses in this strain. Second, use of only female mice limits assessment of sex-dependent innate immune responses to uridine-containing RNA, which is recognized by murine TLR7 and human TLR8. While mouse and human female immune cells show heightened responses to TLR7/8 ligands through TLR7/8 escape from X-chromosome inactivation^73^, downstream murine innate responses to TLR-triggered IL-1 are disproportionately buffered by increased IL-1Ra expression relative to humans^14^. A recent report linking TLR3 genetic polymorphism and mRNA vaccine responses in humans further emphasizes the need to consider species-specific differences when translating our findings^74^. Third, the cellular distribution of STING expression and downstream effects of STING activation differ between species. Human and mouse dendritic cell subsets show distinct STING expression patterns, human STING activation can generate prominent IFN-λ1 responses that are not mirrored in mice, and STING-induced pDC death observed in mice is not recapitulated in human pDCs^75^. Despite these caveats, the use of a defined strain, sex, RNA chemistry, delivery vehicle, and STING agonist enabled controlled dissection of how antigen expression kinetics, formulation-driven biodistribution, and orthogonal STING activation interact.

## Conclusion

This study defines how spatiotemporal colocalization of STING-driven IFN-I signaling and saRNA antigen expression governs the magnitude and polarization of adaptive immunity. By orthogonally engineering adjuvant and antigen biodistribution, we establish that adjuvant and antigen need not be delivered to the same cell to tune vaccine-induced humoral and cellular responses. Synchronizing lymph node delivery of polySTING and LNP-saRNA enhanced the magnitude and polarization of T_H_1 cellular adaptive immunity against both viral and bacterial antigens, although the magnitude of this enhancement varied with prime-boost interval and antigen configuration. These results suggest further opportunities to enhance type I immune responses in T cell-driven immunotherapies. Conversely, mismatching early lymph node IFN-I with later-arriving, muscle-restricted PP antigen expression prolonged saRNA expression at the expense of IFNAR-dependent T cell immunity. This exchange is detrimental to vaccination but may benefit applications in which durable expression and limited T cell reactivity are both desirable, such as protein replacement or tolerizing therapies. These findings provide a blueprint for decoupling innate activation from RNA delivery to rationally shape the adaptive immune response to saRNA vaccines and therapeutics.

## Materials & Methods

### Materials

All chemicals and solvents were purchased from Sigma Aldrich and used without further purification. Phosphate-buffered saline (PBS), RPMI, DMEM, fetal bovine serum (FBS), penicillin-streptomycin (P/S), and HEPES were purchased from Gibco while Normocin®/Zeocin® selective antibiotics, QUANTI-Blue, QUANTI-Luc™, and the mouse macrophage RAW264.7-Dual™ cell line (RAW-Dual) were purchased from Invivogen. Pooled 15-mer peptides with 11 amino acid overlaps from influenza A HA (H1N1 Peptivator) were purchased from Miltenyi Biotech (Germany) while the fLuc-derived peptide^76^ GFQSMYTFV was synthesized by ProImmune (UK) at >80% average purity. Custom peptides 15 amino acids in length with an 11 amino acid overlap spanning amino acids 11-281 of OXA-23 (NCBI Reference: WP_057704766.1) were synthesized at 91.5% average purity by GenScript (China) and resuspended at 20 µg/mL in three sequential pools comprising peptides 1–22, 23–44, and 45–66 for use in ELISpot assays. STING agonist-3 (“di-ABZI”; catalog # HY-103665) and ADU-S100 (“CDN”, a.k.a. ML RR-S2 CDA; catalog # Cat. No.: HY-12885) were purchased from MedChemExpress (USA). Other materials were purchased from suppliers indicated in the text.

### In vitro RNA transcription

Plasmids encoding self-amplifying RNA (saRNA) composed of a Trinidad Donkey (TrD) strain Venezuelan Equine Encephalitis Virus (VEEV) replicase with sub-genomic promoter expression of firefly luciferase (fLuc)^22^, transmembrane Cal/09 H1N1 influenza hemagglutinin (HA_mem_)^22^, and the carbapenem hydrolyzing class D β-lactamase Oxa23 (OXA23)^45^ were cloned and purified in previous work, as was the OXA23 mRNA construct^45^. The transmembrane domain was removed from the HA_mem_ to yield secreted HA saRNA (HA_sec_) using site-directed mutagenesis kit (New England Biolabs) and confirmed by Nanopore sequencing. *In vitro* transcription was performed using 1 μg of linearized DNA template in a HiScribe™ reaction (New England BioLabs, UK) for 2 h at 37 °C, according to the manufacturer’s protocol. Co-transcriptional capping was performed for saRNA constructs with CleanCap AU (TebuBio, UK) or with CleanCap AG (TebuBio, UK) for OXA23 mRNA. Transcripts were then purified by overnight LiCl precipitation at -20 °C, centrifuged at 14,000 RPM for 20 min at 4 °C, washed with 70% EtOH, centrifuged at 14,000 RPM for 10 min at 4 °C, resuspended in nuclease-free water and stored at -80 °C until further use.

### Lipid nanoparticle formulation

Lipid nanoparticle (LNP) formulations were prepared as previously described^44^ Briefly, C12-200, DSPC, plant-derived cholesterol, and DMPE-PEG2000 (Avanti Polar Lipids), and stock solutions were prepared in absolute ethanol. Lipids were mixed at molar ratios of 35:16:46.5:2.5 C12-200:DSPC:Cholesterol:DMPE-PEG2000 at a total lipid concentration of 15 mM. RNA (mRNA or saRNA) solutions were prepared in 50 mM sodium acetate and 100 mM sodium chloride buffer (Ambion), adjusted to pH 5.5. LNPs were formulated using the NanoAssemblr™ Ignite nanoparticle formulation system (Cytiva) at an RNA-to-lipid ratio of 3:1 v/v (1:55 w/w) and a total flow rate of 8 mL/min. Formulated LNPs were diluted in Dulbecco’s phosphate-buffered saline (PBS, Gibco) at a ratio of 1:5 v/v and subsequently concentrated using Amicon Ultra-4 centrifugal filters with a 100 kDa molecular weight cutoff (Merck). The concentration of encapsulated RNA was quantified using the Quant-iT RiboGreen RNA Assay Kit (Thermo Fisher Scientific) according to previously reported methods (routinely > 95%; see source data) and doses specified throughout the text refer to encapsulated RNA. LNP size and polydispersity index (PDI) were determined using a Zetasizer Nano ZS (Malvern Panalytical).

### pABOL synthesis and polyplex formulation

poly(CBA-co-ABOL) “pABOL” was synthesized by aza-Michael polyaddition of 4-amino-1-butanol (ABOL; 1 eq.) to N,N’-cystaminebis(acrylamide) (CBA; 1.004 eq.) reacted for 12 days, purified by dialysis, lyophilized, and analyzed by gel permeation chromatography analysis in LiBr-buffered dimethylformamide according to our published protocol.^22^ The polymer used exhibited molecular weights of M_n_ = 21.0 kDa and M_w_ = 40.7 kDa such that *Ð* = 1.94. Polyplexes (PP) were formulated by separately diluting pABOL to 4.5 mg/mL and saRNA to 0.1 mg/mL (mass ratio 45:1; N/P molar charge ratio of 37:1; previously demonstrated to achieve 100% encapsulation efficiency^22,77^) in equal volumes of 20 mM Bis-Tris pH 6.5 + 5% sucrose w/v (BTS), vortex mixing for 10 s, and diluting further with BTS ± polySTING. PP size and polydispersity index (PDI) were determined using a Zetasizer Nano ZS (Malvern Panalytical).

### PolySTING synthesis and formulation

The polymeric prodrug polySTING was synthesized with and without rhodamine B-methacrylate as in previous work.^43^ The di-ABZI content was quantified by nuclear magnetic resonance spectroscopy (9.3% w/w for polySTING; 8.6% w/w for rhod-polySTING) and used to calculate the polySTING concentration required to obtain the target di-ABZI dosage. For co-administration of polySTING and RNA, polySTING was dissolved at 2X concentration in BTS or PBS and mixed with an equal volume of PP or LNP, respectively, just before injection. For example, a dose of 1 µg PP-saRNA and 10 µg di-ABZI eq. polySTING contains 4.5 µg pABOL and 107.5 µg polySTING polymer in 50 µL of BTS.

### Fluorescence correlation spectroscopy analysis of polySTING-nanoparticle interaction

Fluorescence correlation spectroscopy (FCS) was performed to assess the association of rhod-polySTING with PP- and LNP-saRNA nanoparticles. Rhod-polySTING was mixed with PP or LNP formulations (in BTS or PBS, respectively) at the same dose ratio used for *in vivo* biodistribution studies and incubated for 30 min at room temperature. Samples were diluted 50-fold in their respective formulation buffers immediately prior to measurement to achieve fluorophore concentrations compatible with single-molecule detection. Twenty FCS measurements lasting 10 seconds each were carried out per sample using a confocal Leica SP8 Falcon microscope equipped with a 561 nm laser and avalanche photodiode detector using a 63×/1.2 NA water immersion objective. Autocorrelation curves G(τ) were generated from fluorescence intensity fluctuations and normalized to 1 at τ = 0. The autocorrelation data were fitted using a two-component 3D diffusion model as previously described for protein corona analyses.^78^ The relative amplitude of the slow-diffusing component was interpreted as the fraction of polySTING associated with nanoparticles, while the fast component represented free polySTING in solution, such that a lower bound fraction corresponds to weaker nanoparticle-polySTING interaction. Measurement replicates (n = 20 per sample) were fitted separately using Lecia LAS X software then exported for plotting.

### Mouse transfection/immunization and infection challenge

All animals were handled in accordance with the UK Home OCce Animals Scientific Procedures Act 1986 and with an internal ethics board and UK government approved project (PP5168779; Tregoning) and personal license (I87985646; Peeler). Female BALB/c mice (Charles River, UK) 6−8 weeks of age were placed into groups (N = 5 unless otherwise noted). All cages were kept in a specific pathogen free room at 20-24°C with 55±10% humidity. All immunizations and transfections were 50 µL intramuscular injections into the vastus lateralis muscle per the doses and schedules described in the text. For experiments simultaneously tracking transfection and immunogenicity, fLuc saRNA was injected into the right leg and antigen saRNA was injected into the left leg with equivalent STING agonist doses at both prime and boost timepoints. For infections, mice were anesthetized using isoflurane followed by intranasal instillation of 100 µL PBS containing 3e4 PFU A/California/7/2009 (H1N1) influenza virus^48^ or 5e7 CFU *A. baumannii* BAL_276^45^. Virus was grown in MDCK cells, in serum-free DMEM supplemented with 1 µg/mL trypsin and virus titer was determined by plaque assay. To determine bacterial loads upon challenge, bacteria were serially diluted 1:10 and plated onto agar plates, which were incubated overnight at 37°C to determine the colony forming units per mL.

### Depletion studies

Neutrophils were depleted by intraperitoneal injection of 500 µg anti-mouse Ly6G (clone 1A8, rat IgG2a κ, *InVivo*Plus, Bio X Cell BP0075-1, RRID AB_2894803) or an equal dose of isotype-matched control (anti-trinitrophenol, clone 2A3, rat IgG2a κ, *InVivo*Plus, Bio X Cell BP0089, RRID AB_1107769) diluted in <500 µL sterile PBS immediately before use. Antibody was dosed one day before and one day after each immunization for a total of four doses. Mice were primed with 1 µg fLuc/HA_mem_ saRNA ± 10 µg polySTING on day 0, bled and boosted with the same formulations 4 weeks later, and sacrificed at week 6 for serum antibody and splenocyte analysis. Luminescence was imaged on days 3, 7, 14, and 21.

### IFNAR blockade

Type I interferon signaling was blocked 24 h before each immunization by intraperitoneal injection of 1 mg anti-mouse IFNAR-1 (clone MAR1-5A3, mouse IgG1, low endotoxin, Assay Genie IVMB0202) diluted in <500 µL sterile PBS, as previously described^48^. Mice were primed with 1 µg HA_mem_ saRNA on day 0 and bled 6 h later for serum cytokine analysis. Blood was collected 48 h before the boost at 4 weeks to assess post-prime titers and sacrifice was at week 6 for serum and spleen collection.

### Serum cytokine quantification

Serum was isolated as the supernatant following centrifugation (15,000 g, 15 minutes) of blood collected through venipuncture and stored at -80 °C until assay. Cytokines in blood were assessed using a custom mouse kit from Meso Scale Discovery (MSD) as a 10-spot U-PLEX kit (K15069L-2), including the following analytes (lower limit of detection in pg/mL in parentheses): IFN-α2 (30.7), IFN-β, (1.7), IFN-g (1.0), IL-12p70 (35.2), IL-6 (5.7), CXCL1 (0.7), CXCL10 (1.5), CCL2 (0.4), CCL3 (0.7), and CCL4 (8.9). Sera were diluted 0.2-0.4-fold depending on the volume collected; mice were excluded from analysis when serum yield was insufficient for analysis.

### Longitudinal luciferase expression

At each timepoint indicated, mice were anesthetized with isoflurane for *in vivo* fLuc bioluminescence imaging in the Spectral Instruments AMI ten minutes after intraperitoneal injection of 100 µL of VivoGlo™ Luciferin (ProMega). Imaging data were analysed in Spectral Instruments Aura software (version 4.0.7) to report the total emission from a uniform region of interest in terms of photons/sec.

### Flow cytometry

Cell suspension preparation and staining followed previously published work^24,48^. Mouse quadriceps were digested in 1 mL Liberace (12.5 mg/mL, Roche), DNase (200 mg/mL, Sigma), and hyaluronidase (50 mg/mL, Molecular Dimensions) in RPMI 1640 medium with 10% FBS + 1X P/S + 10 mM HEPES (cRPMI) with shaking for 1 h at 37 °C, while lymph nodes were digested in DNase (200 mg/mL) in cRPMI for 15 min. Following digestion, tissues were forced through 100 µm cell strainers and centrifuged at 500 g for 5 min. Cell pellets were treated with red blood cell lysis buffer (ACK; 0.15 M ammonium chloride, 1 M potassium hydrogen carbonate, and 0.01 mM EDTA, pH 7.2) before centrifugation at 200 g for 5 min. Cells were resuspended in cRPMI, and viable cell numbers determined by trypan blue exclusion. Cells were incubated in a U-bottom 96-well plate for 20 min at 4 °C in the dark with 100 µL Live/Dead violet dye (Thermo Fisher Scientific, L34955), then centrifuged at 500 g for 2 min and the supernatant was discarded. The cell pellet was resuspended in Fc block (BD cat. # 6266549) in PBS + 1% BSA (PBSA) and stained with antibodies for the following surface markers: CD3-APC (Clone: 17A2, Invitrogen, Cat. #17-0032-82), Ly6G-BV605 (Clone: 1A8, BD, Cat. #563005), CD11c-PE (Clone: HL3, BD, Cat. #557401), F4/80-SB780 (Clone: BM8, eBiosciences, Cat. #78-4801-82), CD103-PerCP-eFluor710 (Clone: 2E7, eBiosciences, Cat. # 46-1031-82), NK1.1-APC-Cy7 (Clone: PK136, BD Biosciences, Cat. #560618), and MHCII-eFluor450 (Clone: M5/114.15.2, eBiosciences, Cat. # 48-5321-82) for 1 h at room temperature in the dark. Cells were washed with PBSA three times before filtration into FACS tubes. Cells were acquired on an LSR Fortessa Flow cytometer (BD Biosciences) and analyzed with FlowJo software (BD Biosciences; v11.0). Fluorescence minus one controls were used for surface stains. The gating strategy is shown in Fig S3.

### Confocal microscopy

To assess the spatial distribution of saRNA expression *in vivo*, Ai14 reporter mice were transfected IM with 10 µg (di-ABZI eq.) polySTING ± 5 µg Cre saRNA in PP or LNP. Quadriceps muscles and inguinal lymph nodes were harvested 24 h post-transfection, fixed in 4% paraformaldehyde (PFA) in PBS, and processed either for whole-tissue clearing followed by spinning-disk confocal microscopy or for cryosectioning and laser-scanning confocal microscopy.

For whole-mount spinning disk confocal imaging, tissues were immunostained using a modified 2Eci clearing protocol at room temperature^79^. Briefly, samples were permeabilized and blocked for one day in a solution containing 10% 10×PBS, 2% Triton X-100, 20% DMSO, 5% BSA (PBS-TxDB; all v/v % in MilliQ), followed by six days of incubation with primary antibodies against tdTomato (rabbit monoclonal, clone RF5R; Thermo Fisher Scientific) and Ly6G (rat monoclonal, clone RB6-8C5; Thermo Fisher Scientific) at 1:50 dilution in PBS-TxDB. Following three days of washing with PBS-TxDB, tissues were incubated for five days with donkey anti-rabbit-AlexaFluor555 and anti-rat-AlexaFluor488 secondary antibodies (Invitrogen) at 1:50 dilution in PBS-TxDB. Secondary antibody staining as well as rinsing/washing procedure were the same as for primary antibodies. After washing, tissues were transferred into PBS, washed for >1h and subsequently fixed with 4% PFA for 4 hours. Tissue was then stored in PBS until ready for tissue clearing and imaging. For tissue clearing, tissues were transferred into 45ml 30%, 50%, 70% and 2x 100% MeOH (Sigma, Cat.No. 5895960100) for 1 day per step on a tube roller. Notably, anhydrous MeOH was used for the last 2 100% MeOH steps. Tissue was then transferred into stabilized Ethyl Cinnamate (Sigma,Cat.No. W243000). Microscopy was performed using a Nikon Spinning Disk Confocal (Nikon Eclipse Ti2 with a Crest X-Light V3 Spinning Disk and Ultra-Long Working Distance Objectives using a custom printed imaging plate.

Lymph node volume was calculated from serial cross-sectional area measurements along the z-axis using trapezoidal numerical integration. Z distance was corrected by Refractive Index (Ethyl Cinnamate)/RI(Air) (thus, z*1.56= true z distance). For each sample, the projected area of the region of interest was measured at discrete z-positions (100µm steps). The volume was then estimated by summing the volumes of adjacent z-segments.

For standard confocal imaging, fixed tissues were embedded in Optimal Cutting Temperature (OCT) liquid, frozen on dry, and sectioned into 10 µm-thick sections on a microtome. Sections were incubated with 1 µM DAPI in PBS and imaged using a Leica SP8 Falcon equipped with a 60X 1.3 NA water immersion objective. Confocal z-stacks were acquired at 0.5 µm spacing and processed using Fiji (version 2.9.0), where images were converted into 5 µm-thick maximum intensity z-projections.

### Serum antibody quantification by ELISA

Antigen-specific IgG antibody titers in serum were measured using an endpoint ELISA. High-binding Costar plates (Corning) were coated with recombinant HA at 1 μg/mL in PBS^22^ or *A. baumannii* BAL_276 whole bacterial cells resuspended at OD_600_ 0.5 in carbonate buffer^45^ (for samples) or anti-mouse IgG capture antibodies at 5 µg/mL (for standards; Southern Biotech) in PBS at 4 °C overnight. Highly purified polyclonal mouse IgG, IgG1, or IgG2a (Southern Biotech) were used to generate a standard curve in a 1:5 dilution series, starting at 200 ng/ml. Plates were washed three times (PBS + 0.1% Tween-20 w/v) and then blocked with assay buffer (PBS + 1% w/v BSA) for 1 h at 37 °C. Serum samples were serially diluted in assay buffer and incubated on the coated plates for 2 h at 37 °C. Plates were washed three times and HRP-conjugated donkey-anti-mouse IgG, IgG1, or IgG2a (1:5000, Southern Biotech) was added for 1 h at 37 °C. Plates were washed three times, developed with TMB Substrate (Thermo Fisher) and stopped with 2 N H_2_SO_4_ (Sigma). Absorbance at 450 nm was measured using a SpectraMax plate reader. Titers were interpolated from background-subtracted absorbance values using a sigmoidal, four point least squares fit of the standard curve (GraphPad Prism) and scaled by the dilution factor used.

### Splenocyte ELISpot

Spleens were mashed through 70 µm cell strainers to yield splenocyte suspensions, washed with cRPMI, treated with ACK lysis buffer, counted, and diluted to 1e7 cell/mL. Splenocytes were stimulated with 1.25 µg/mL anti-CD28 (Clone 37.51, BD) and either 1 µg/mL HA peptide pool, luciferase peptide, or OXA23 peptide pool (see *Materials*) overnight in cRPMI in the IFN-C ELISpot assay plate (AbCam ab64029) or the FluoroSpot Plus: Mouse IFNCγ/ILC17A assay plate (Mabtech FSP-3144-2) and then developed according to the manufacturer’s protocol. The spots were counted using the AID iSpot reader and EliSpot Reader software V7.0.

### Influenza viral load

Viral load was assessed by Qiazol extraction of RNA from frozen lung tissue disrupted in a TissueLyzer (Qiagen). RNA (200 ng) was reverse-transcribed into cDNA using random hexamer primers (ProMega GoScript) and quantitative PCR of the influenza M gene was carried out using 2X TaqMan Universal Master Mix II (ThermoFisher), 0.1 µM forward primer (5’-AAGACAAGACCAATYCTGTCACCTCT-3’), 0.1 µM reverse primer (5’-TCTACGYTGCAGTCCYCGCT-3’), and 0.2 µM probe (5’-FAM-TYACGCTCACCGTGCCCAGTG-TAMRA-3’) on a Stratagene Mx3005p system (Agilent Technologies). An influenza M gene plasmid was used as standard to determine M-specific RNA copy number^48^.

### Quantification of bacterial load

Lungs and spleens were homogenized using the gentleMACS tissue dissociator (Miltenyi). Organ homogenates were then serially diluted 1:10 and plated onto HiChrome Acinetobacter Agar Base (HiMedia Labs) plates and incubated overnight at 37 °C to yield countable spots^45^.

### Cell culture, transfection, and reporter assays

RAW-Dual cells were cultured in complete DMEM supplemented with 10% FBS, 1X P/S, and 1X Normocin®/Zeocin® in a humidified incubator at 37 °C with 5% CO_2_ and regularly tested negative for mycoplasma contamination using a DNA-based PCR test. Cells were seeded overnight in a 96-well plate at a density of 1e5/100 µL media per well and exchanged to 200 µL fresh media prior to transfection with 150 ng of PP or LNP formulated saRNA with or without the doses of STING agonists mentioned in the text. Twenty-four hours later, the media was removed and assayed for IRF and NF-κB activation using QUANTI-Luc™ and QUANTI-Blue reagents, respectively, while cells were assayed for saRNA transfection using the ONE-Glo firefly luciferase assay system (ProMega) according to the manufacturers’ instructions. Absorbance (NF-κB) and luminescence (IRF, fLuc) were measured using an EnVision plate reader (Revvity) and data from untreated cell controls was used for background subtraction.

### Statistical Analysis

Datasets that are log-distributed by measurement scale (transfection efficiency, *in vivo* fluorescence efficiency, endpoint ELISA titers, IgG subclass ratios, bacterial colony-forming units, and viral genome copy number) were log10-transformed prior to analysis, plotted on log10 axes, and are summarized as the geometric mean with 95% confidence interval. All other datasets (relative body weight, ELISpot spot-forming units, flow cytometry cell counts, hydrodynamic diameter, luminescence area under the curve, and serum cytokine concentration) were analyzed on untransformed values, plotted on linear axes, and are summarized as the mean ± standard deviation. Unless otherwise indicated in the figure legend, groups were compared by one-way or two-way ANOVA with Tukey’s multiple-comparisons correction applied to the values as transformed above. Where cytokine concentrations are displayed as heatmaps spanning multiple analytes of differing dynamic range, a logarithmic color scale is used for legibility only and statistical comparisons were performed on the measured values. Formal normality testing was not performed given the group sizes used here (N = 3 to 8). The transformation applied to each dataset was instead determined by the measurement scale of the underlying assay, and ANOVA was selected for its robustness to moderate departures from normality when group sizes are comparable.

Where limited serum volume produced unequal group sizes with values near the assay detection limit (Fig 1f, Fig S2, Fig S3), and where censoring at the limit of detection (LOD) produced groups with negligible variance (Fig 2, Fig 5e-f, Fig 6m), groups were compared by Kruskal-Wallis test with Dunn’s multiple-comparisons correction. Values below the LOD were imputed on the LOD as indicated by a dashed line. Where the quantity of interest was the fold change between paired treatment groups rather than the absolute concentration (Fig 8b), analysis was performed on log2-transformed concentrations so that increases and decreases of equal magnitude are treated symmetrically. Statistical analysis and curve fitting were performed in Prism 10.3.1 (GraphPad Software) and *p* < 0.05 was considered significant.

## Data availability statement

All the data supporting the results in this study are available within the paper and its Supplementary Information. Raw source data are provided with this paper.

## Acknowledgments

We thank Lesley Rawlinson for lab management support and KS for *in vivo* study support, as well as Professor Simone Di Giovanni and Hee Hwan Park for the kind donation of the Ai14 mice used in this study. D.J.P., J.T., and Z.W. acknowledge support by the BactiVac Catalyst award number BVNCP8-03 through the UK MRC, the International Science Partnerships Fund, and The Department of Health and Social Care as part of the Global AMR Innovation Fund. JT and LS were supported by MRC grant UKRI5802. The views expressed in this publication are those of the authors and not necessarily those of the UK Department of Health and Social Care. D.J.P. was supported by the European Union’s Horizon 2020 research and innovation programme under Marie Skłodowska-Curie grant agreement 101027174. P.S.S. acknowledges support from the U.S. National Institutes of Health, National Cancer Institute grant R01CA257563 (P.S.S.) and the U.S. National Institutes of Health National Institute of Allergy & Infectious Diseases grant R01AI134729. M.M.S. acknowledges support from the Department for Science, Innovation and Technology and the Royal Academy of Engineering Chair in Emerging Technologies award (CiET2021\94), the Rosetrees Trust, and the University of Oxford Strategic Research Fund. R.J.S. acknowledges support from the Department of Health and Social Care via UK Aid funding managed by EPSRC through grant numbers EP/Y530529/1 and EP/Y010167/1 as well as EPSRC and MRC through UKRI IAA award numbers EP/X52556X/1 and MR/X502959/1.

## Author contributions

D.J.P. led the conceptualization of the study, with support from P.S.S., R.J.S., and J.S.T. Methodology was developed by D.J.P., Z.W., C.T., L.S., D.R., K.S., P.F.M., D.C.N., P.S.S., R.J.S., and J.S.T. Investigation was performed by D.J.P., Z.W., S.J., C.T., L.S., D.R., K.S., P.F.M., D.C.N., and J.S.T. Formal analysis was carried out by D.J.P., Z.W., L.S., D.R., and J.S.T. Resources were provided by P.S.S., M.M.S., R.J.S., and J.S.T. Visualization was performed by D.J.P. and D.R. D.J.P. and J.S.T. wrote the original draft of the manuscript and all others contributed to writing, review, and editing of the manuscript. Supervision was provided by D.J.P., M.M.S., R.J.S., and J.S.T. Funding was acquired by D.J.P., P.S.S., M.M.S., R.J.S., and J.S.T.

