## Supplementary Information for "Spatiotemporal control of STING activation and saRNA delivery decouples humoral and cellular immunity"

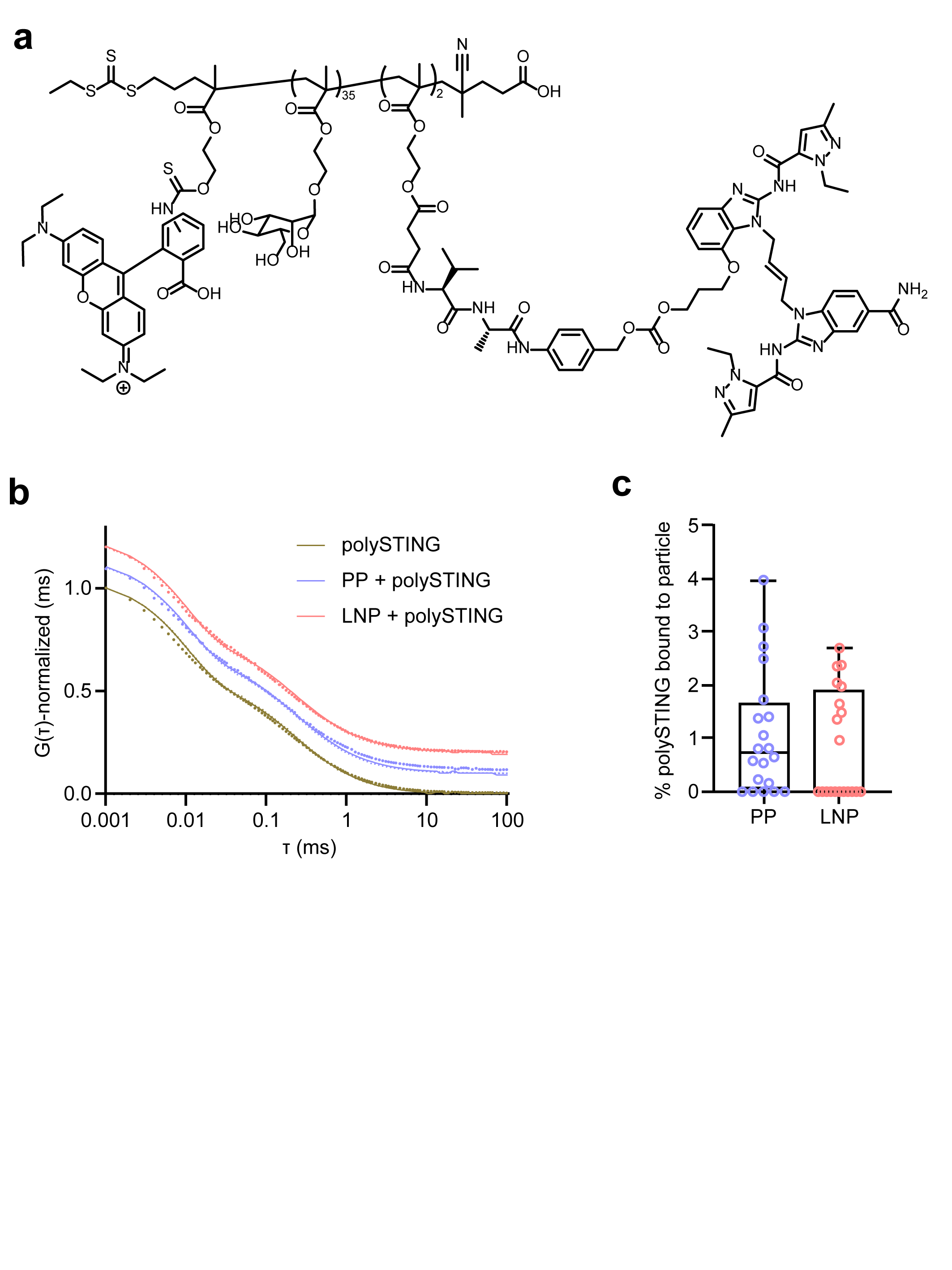

**Figure S1: Fluorescence correlation spectroscopy (FCS) analysis of polySTING association with PP and LNP.** **(a)** Chemical structure of rhodamine-polySTING. Rhodamine-polySTING was mixed with buffer, PP, or LNP formulations at the same concentrations used for *in vivo* administration, incubated for 30 min at room temperature, and diluted 50-fold in PBS prior to FCS measurement. Autocorrelation functions G(τ) from n = 25 technical repeats of N = 1 independent experiment were recorded and analyzed by a two-component diffusion model to quantify the fraction of polySTING associated with nanoparticles. **(b)** Normalized autocorrelation curves (G(τ), offset for clarity) with mean experimental data shown as dotted lines and two-component fits as solid lines. **(c)** Percentage of polySTING bound to nanoparticles as determined from the slow-diffusing component of the two-component fit. Box plots display the median as the central line, the interquartile range as the box boundaries, and the minimum and maximum values as the whiskers.

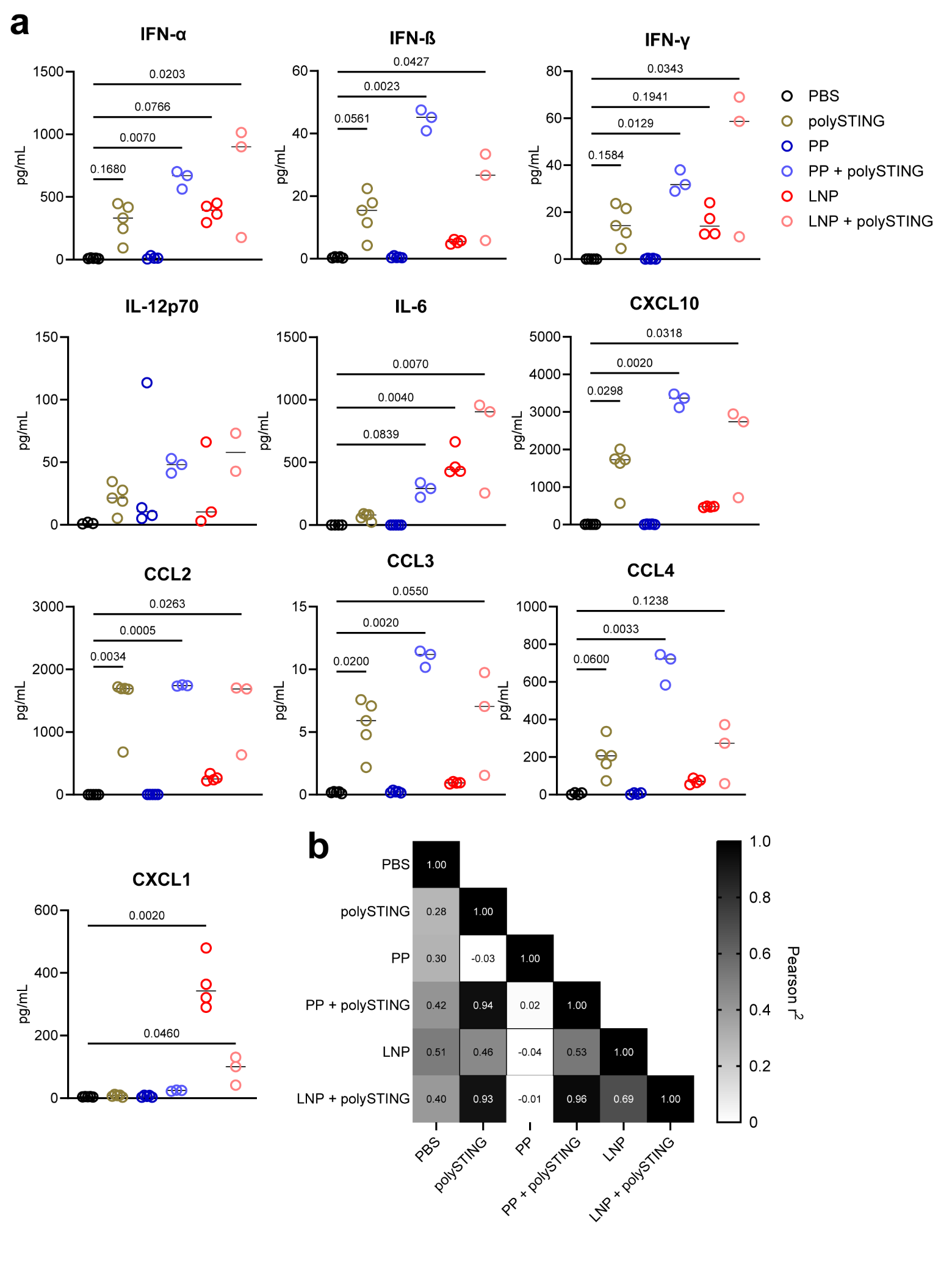

**Figure S2: Acute cytokine responses and correlation analysis.** **(a)** Serum cytokine concentration 6 h after IM 1 µg saRNA ± 10 µg polySTING. *N*=3-5; samples with insufficient serum volume were excluded. Significance calculated via Kruskal-Wallis test with Dunn’s multiple-comparisons correction on raw (non-log-transformed) data. **(b)** Heatmap of Pearson’s correlation matrix of treatment-induced cytokine responses.

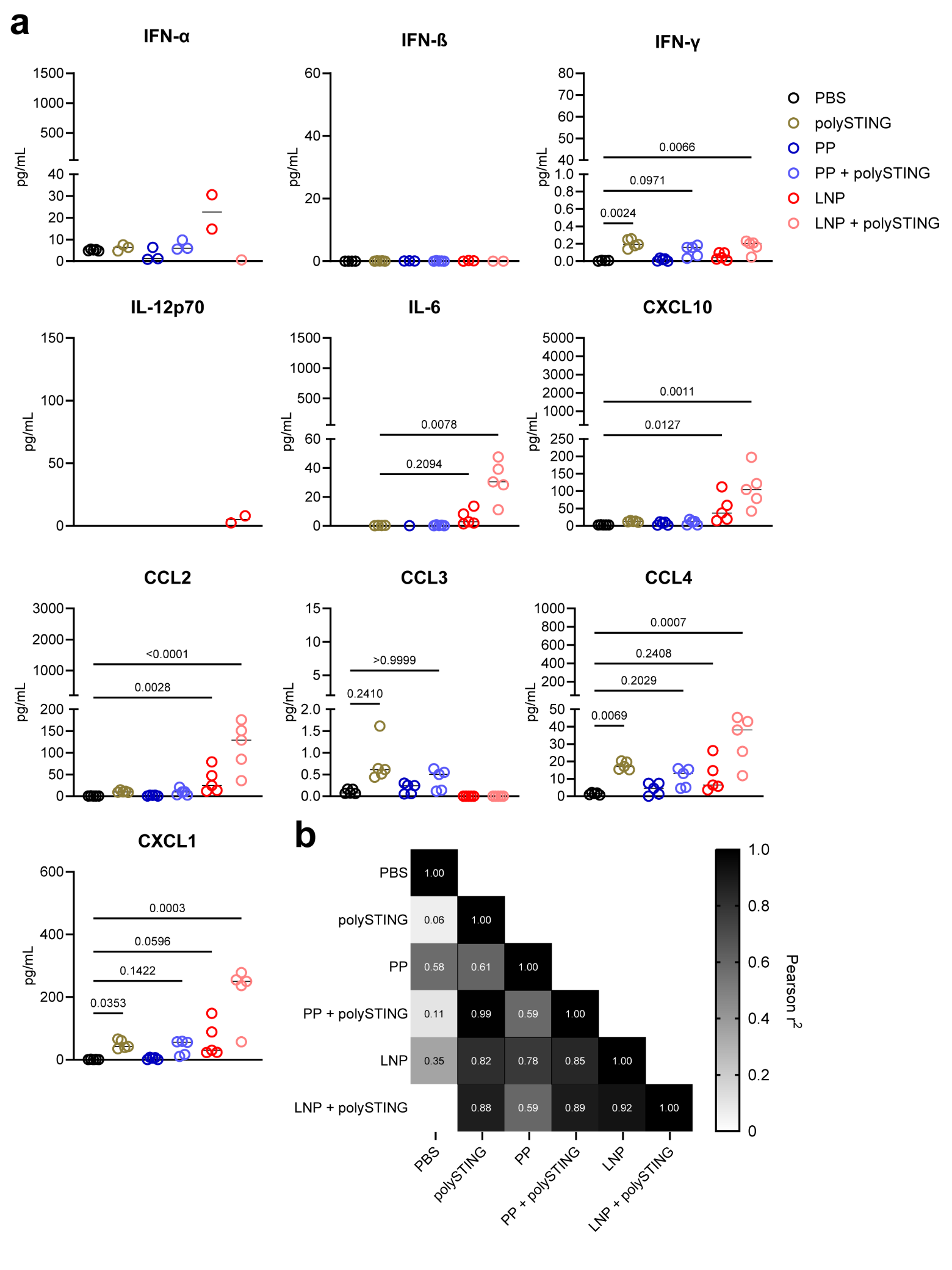
**Figure S3: Resolution of acute cytokine responses.** **(a)** Serum cytokine concentration 48 h after IM 1 µg saRNA ± 10 µg polySTING. *N*=3-5; samples with insufficient serum volume were excluded. Significance calculated via Kruskal-Wallis test with Dunn’s multiple-comparisons correction on raw (non-log-transformed) data. **(b)** Heatmap of Pearson’s correlation matrix of treatment-induced cytokine responses.

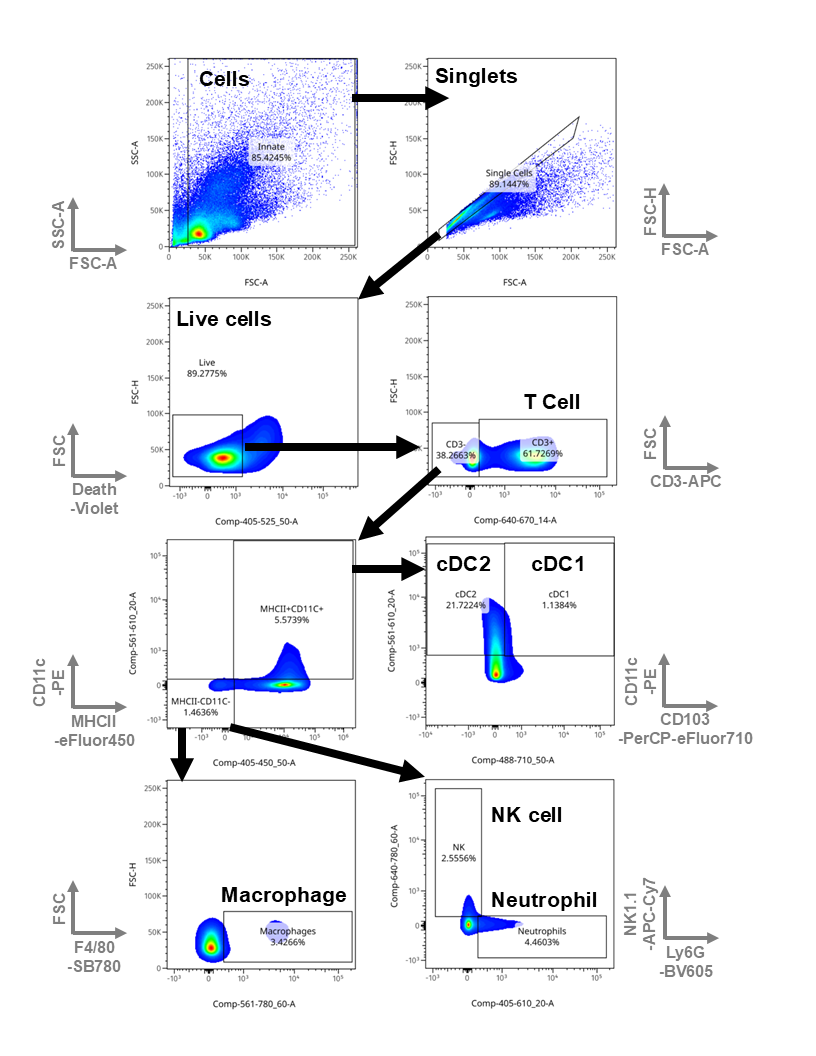

**Figure S4: Innate cell flow cytometry gating strategy.** Cells were extracted from lymph nodes or muscle prior to running by flow cytometry. Gating strategy for different cell types shown.

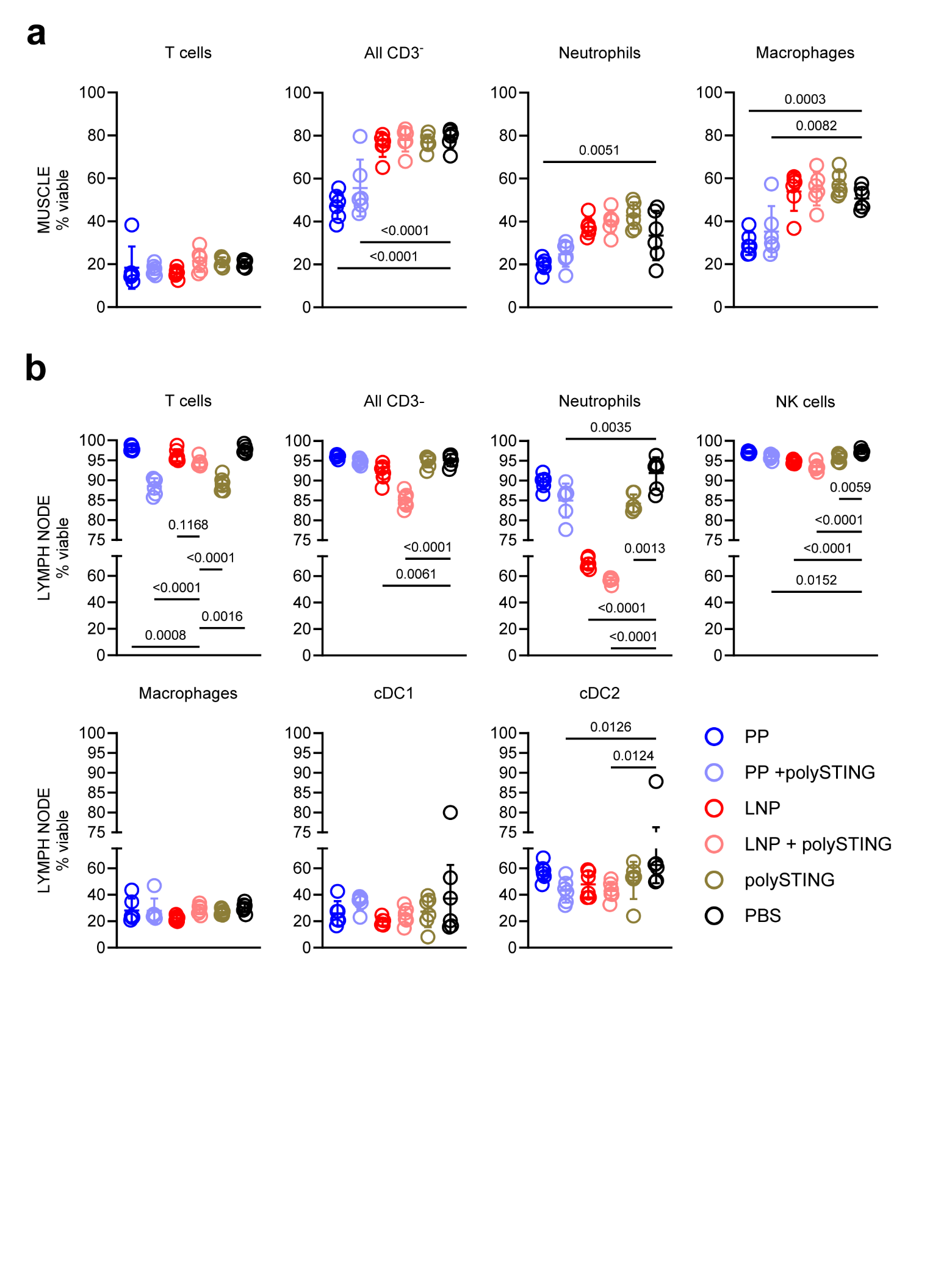

**Figure S5:** **Flow cytometry analysis of cell viability.** Mouse quadriceps muscle and inguinal lymph nodes (*N* = 6) were analyzed by flow cytometry 24 h after IM 10 µg mGL saRNA ± 10 µg (di-ABZI eq.) rhodamine-labelled polySTING. Frequencies of viable singlets in **(a)** muscle and **(b)** lymph node are shown as the mean ± SD, with significance determined by one-way ANVA with Tukey’s HSD test.

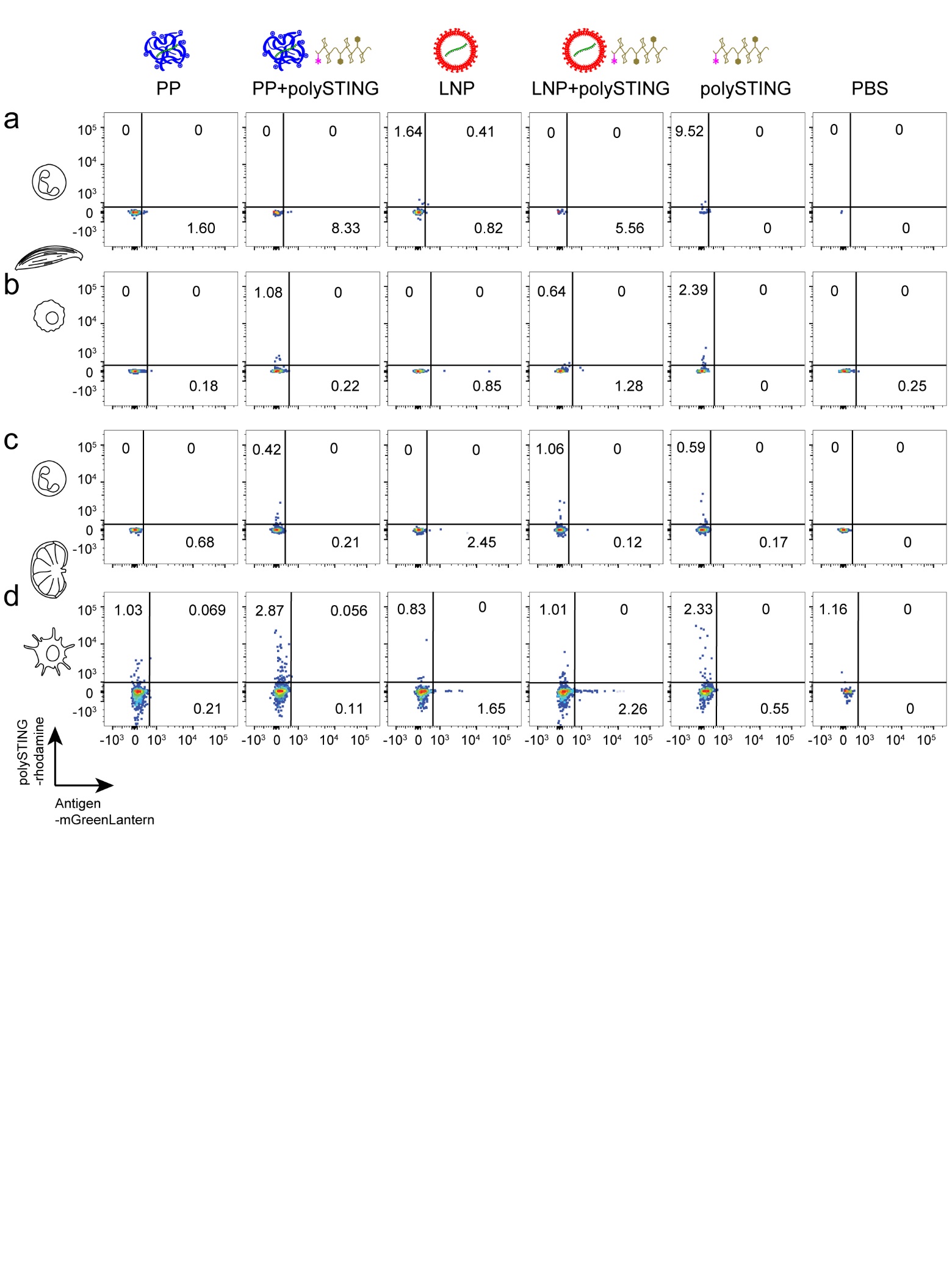
**Figure S6: Antigen and adjuvant flow cytometry gating strategy.** Gating strategy shown for the major cell types taking up antigen (mGreenLantern) and adjuvant (rhodamine-labeled polySTING) in the muscle **(a)** neutrophils and **(b)** macrophages as well as lymph node **(c)** neutrophils and **(d)** cDC2.

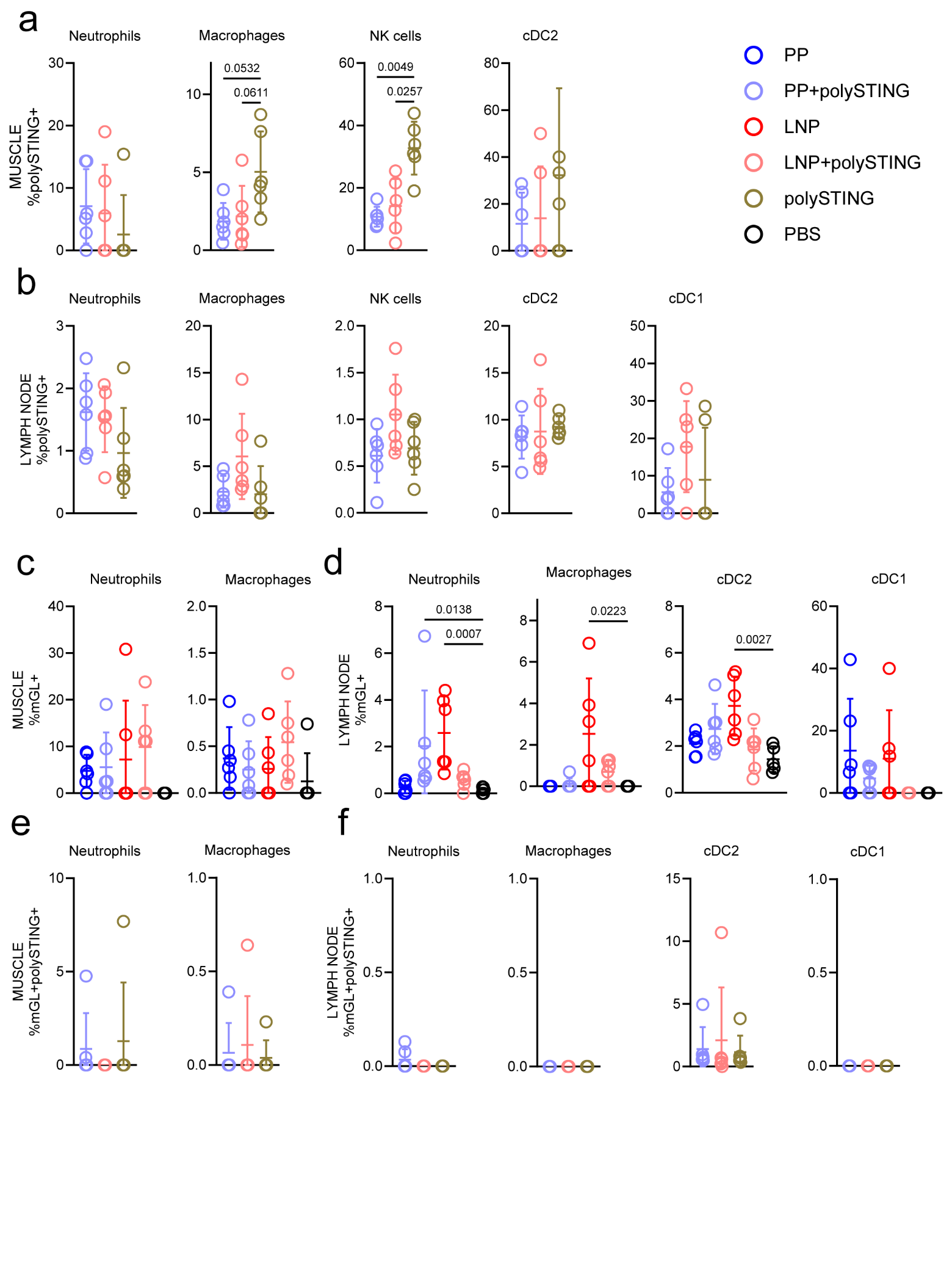

**Figure S7: Flow cytometry analysis of polySTING^+^ and mGL^+^ cell frequencies.** Mouse quadriceps muscle and inguinal lymph nodes (*N* = 6) were analyzed by flow cytometry 24 h after IM 10 µg mGL saRNA ± 10 µg (di-ABZI eq.) rhodamine-labelled polySTING. Shown are cell-type specific rhodamine-polySTING^+^ viable singlets in **(a)** muscle and **(b)** lymph node; cell-type specific mGL+ viable singlets in **(c)** muscle and **(d)** lymph node; and cell-type specific rhodamine-polySTING^+^/mGL+ viable singlets in **(e)** muscle and **(f)** lymph node. Significance determined by Kruskal-Wallis test with Dunn’s correction (mean ± SD shown).

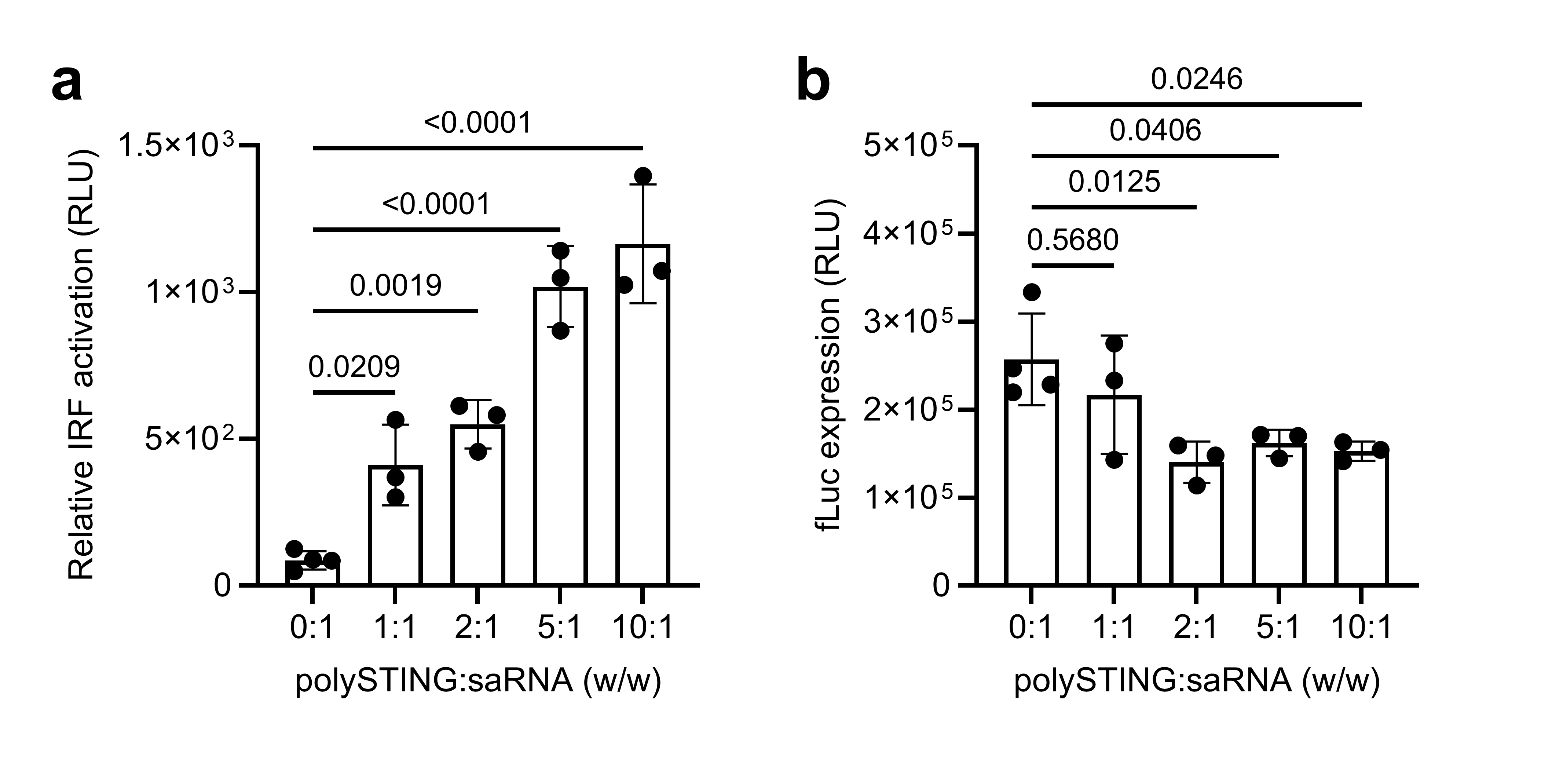

**Figure S8: Colocalized polySTING and LNP delivery increases innate activation and decreases saRNA expression *in vitro*.** Various di-ABZI eq. doses of polySTING were mixed with LNP delivering 150 ng fLuc saRNA to cultured RAW 264.7 Dual macrophages (*N*=3-4). After 24 h incubation, **(a)** activation of gaussian luciferase-linked interferon response factor (IRF) and **(b)** firefly luciferase expression were analyzed through gLuc and fLuc bioluminescence assays, respectively (mean ± SD shown). The average signal from untreated cells was subtracted from all data.

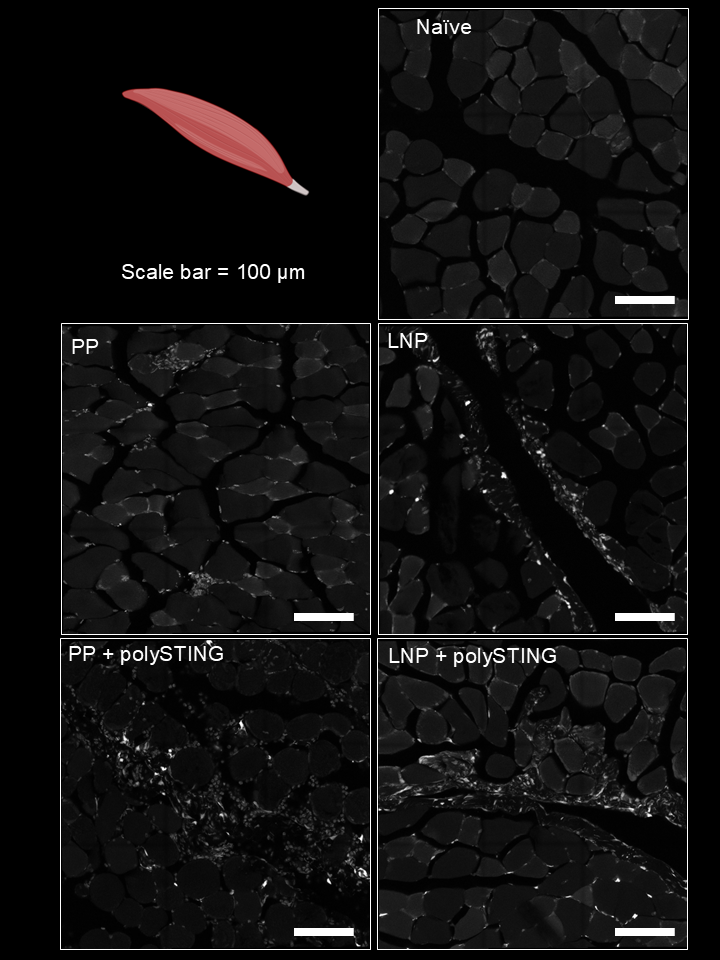
**a**

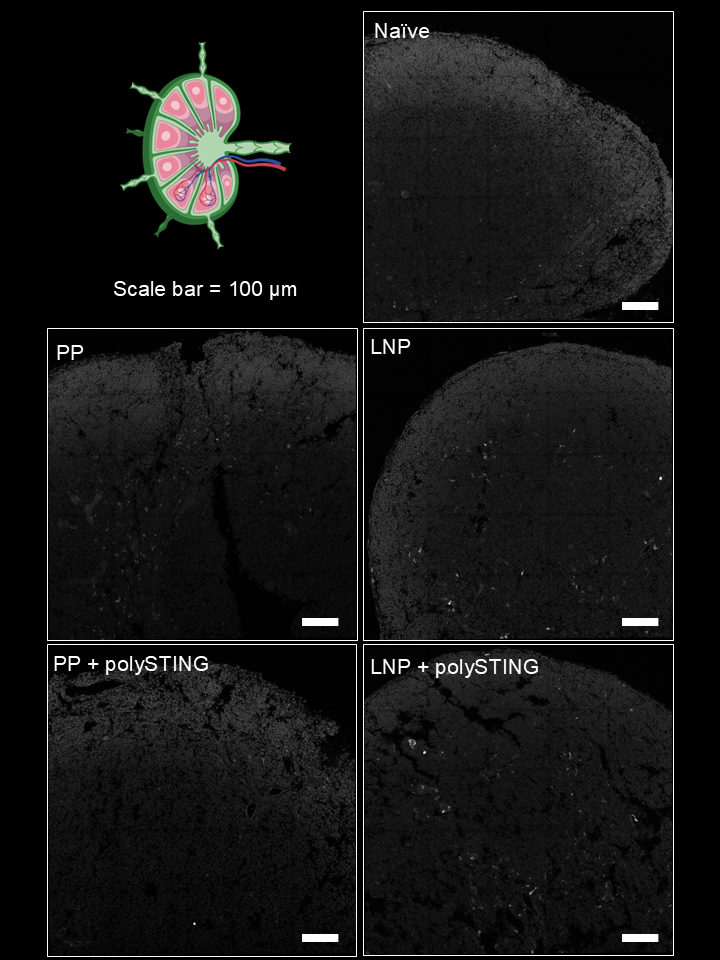
**b**

**Figure S9: Confocal micrographs of tdTomato expression in Ai14 mice.** Female Ai14 mice (N = 1 mouse per group; n = 2 injections per mouse) were injected intramuscularly bilaterally with either PBS or 5 µg Cre saRNA ± 10 µg (di-ABZI eq.) polySTING. The **(a)** quadriceps and **(b)** inguinal lymph nodes were dissected 24 h later, fixed, and either tissue cleared for spinning disk confocal microscopy (shown in **Fig 4**) or cryo-sectioned for laser scanning confocal microscopy. Images were converted to 5 µm-thick maximum intensity z-stacks using ImageJ software to reveal transfected (tdTomato^+^) cells.

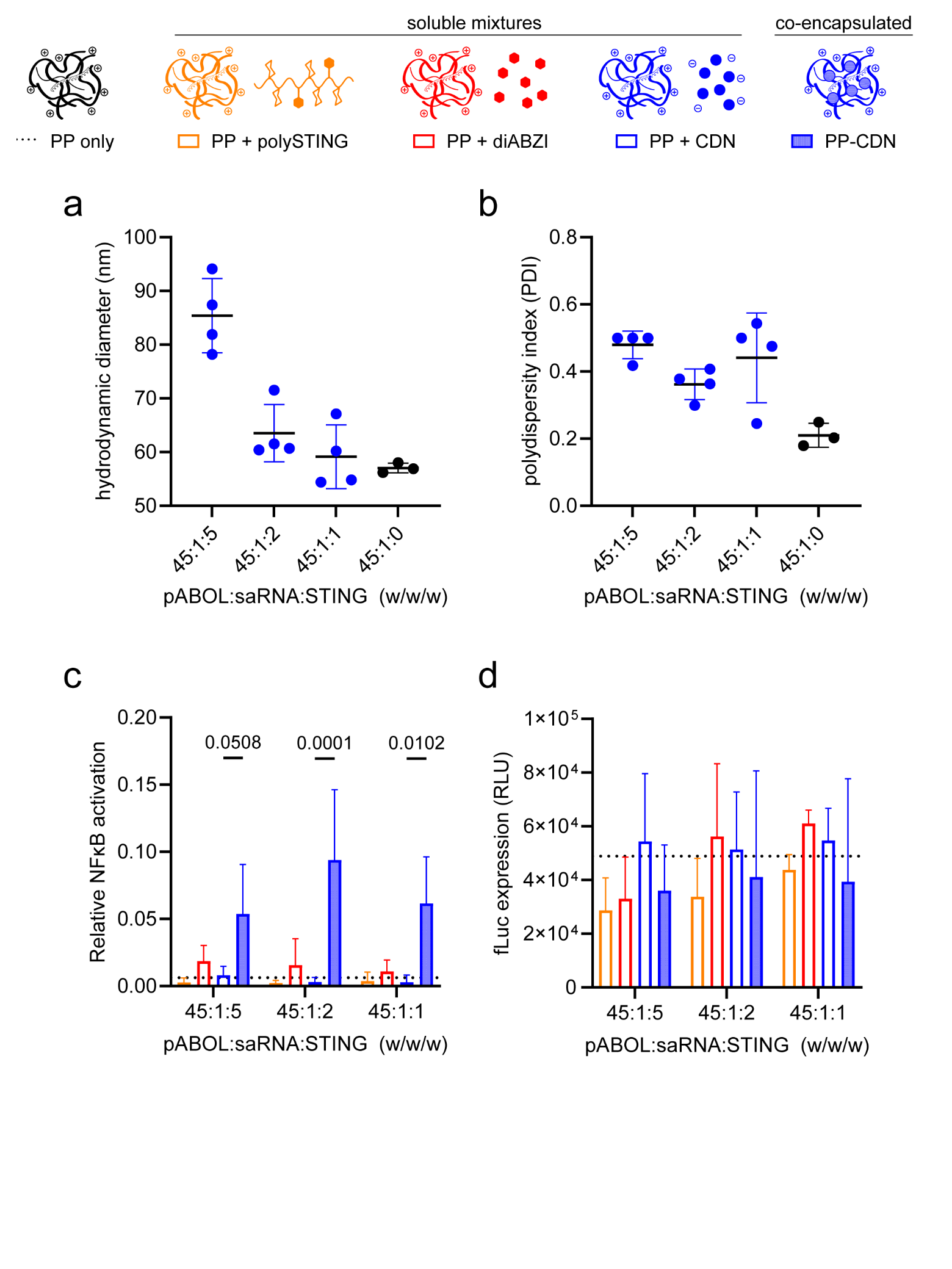

**Figure S10: PP can encapsulate both saRNA and various concentrations of a cyclic dinucleotide STING agonist (ML-RR-CDA; “CDN”).** Various weight/weight ratios of CDN and fLuc saRNA were co-encapsulated by pABOL to form polyplexes and measured by dynamic light scattering. **(a)** Hydrodynamic diameter and **(b)** polydispersity index data of *N* = 3-4 separately formulated PP batches shown as mean ± SD. Various STING agonists were either mixed with (di-ABZI, polySTING, CDN) or co-encapsulated in (PP-CDN) polyplexes. After 24 h incubation of RAW 264.7 Dual macrophages with 150 ng fLuc saRNA ± various relative STING agonist doses (*N* = 3), activation of **(c)** nuclear factor κB (NF-κB) and **(d)** luciferase expression were analyzed. Dashed line shows the mean results of PP only. Significance determined by one-way ANOVA with Tukey’s HSD test (mean ± SD shown). The average signal from untreated cells was subtracted from all data.

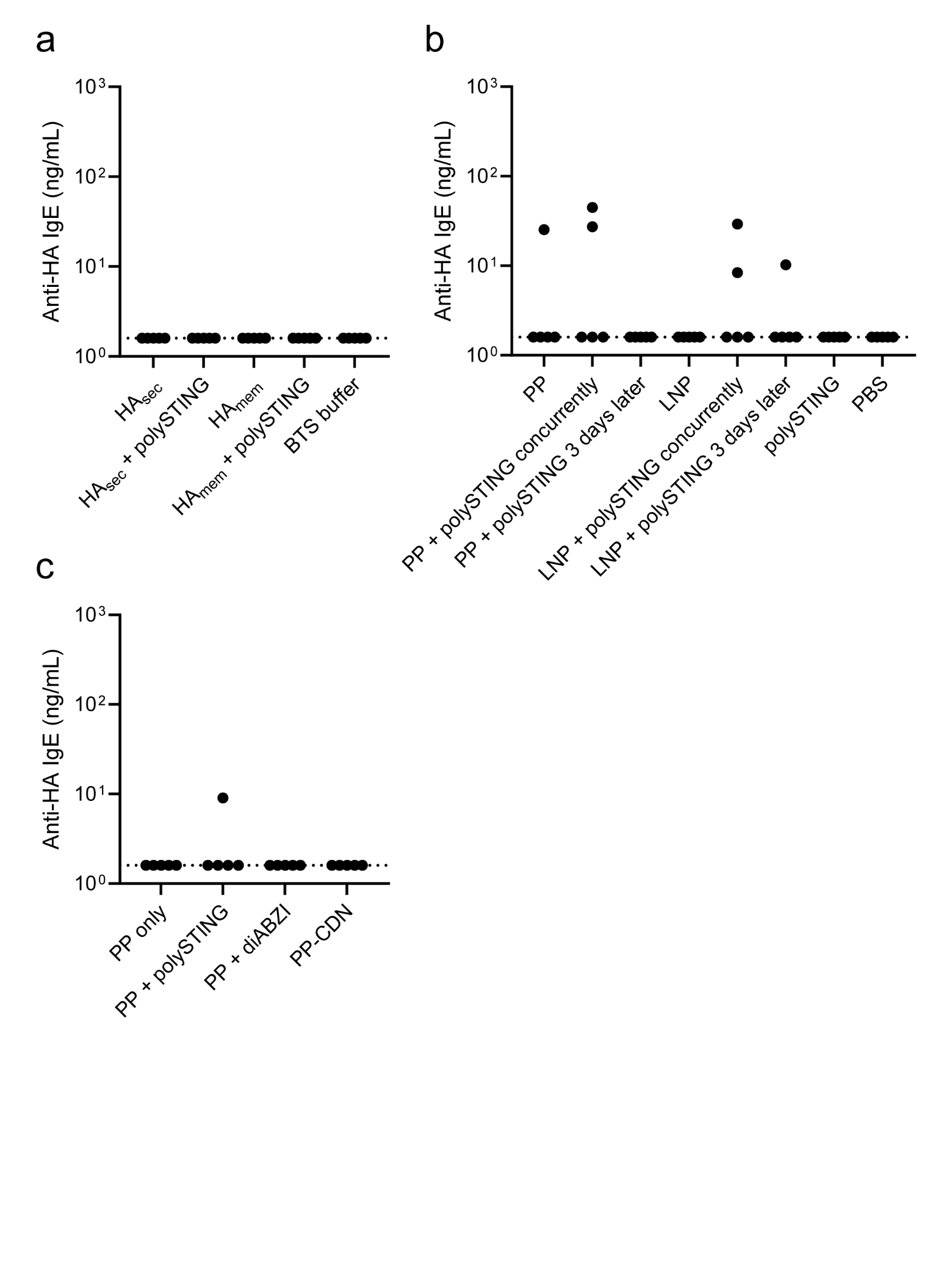

**Figure S11: Evaluation of IgE immune responses.** Serum collected at harvest from mice in the studies described in (**a-b**) Figures 6 and (**c**) Figure 7 were analyzed by ELISA for HA-specific IgE titers using a minimal 1:10 dilution in assay buffer. Signals below assay limit of detection were plotted on the dotted line (N =5).

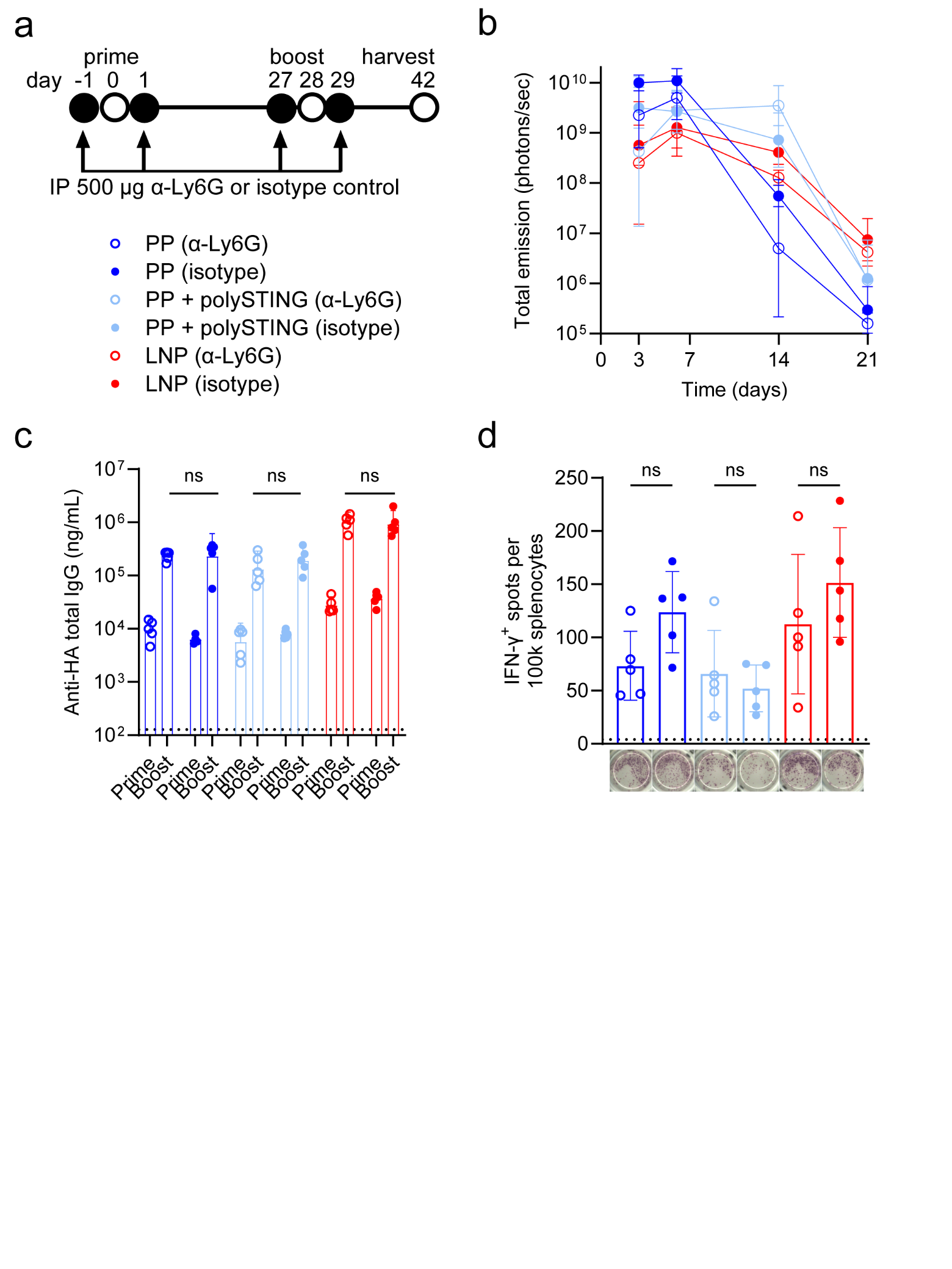

**Figure S12: The role of neutrophils in polySTING-modulated transfection and immunogenicity. (a)** Mice were dosed intraperitoneally (IP) with anti-Ly6G or isotype-matched antibodies on the day before and after receiving 1 µg fLuc (right leg) and HA (left leg) saRNA ± 10 µg polySTING according to the prime/boost schedule shown (*N* = 5). **(b)** Luciferase expression was analyzed by *in vivo* imaging. HA-specific adaptive responses were quantified by **(c)** serum IgG ELISA and **(d)** splenocyte IFN-ɣ ELISpot (pictures of representative wells shown below).

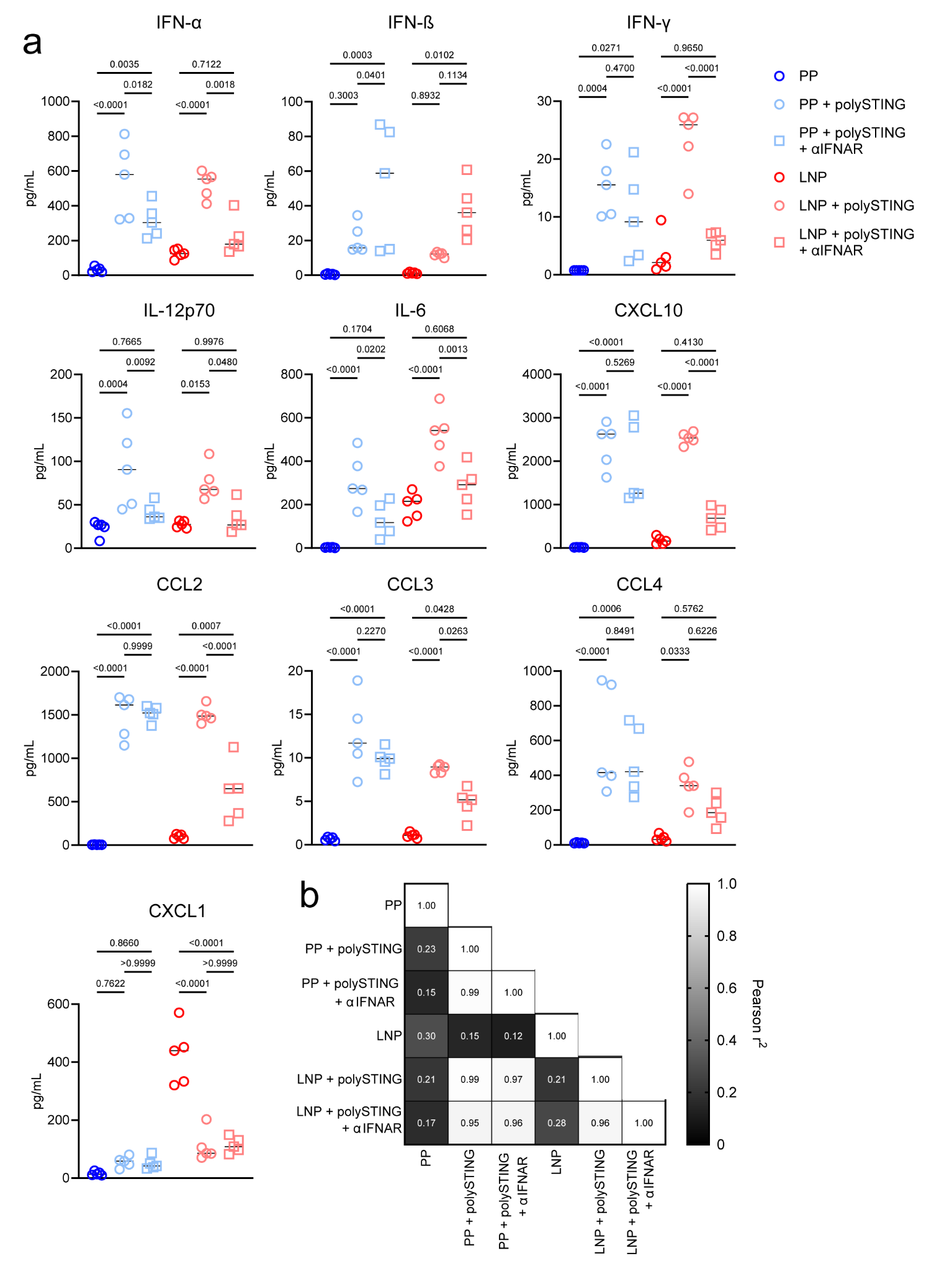

**Figure S13: Acute cytokine responses under IFNAR blockade.** **(a)** Serum cytokine concentration 6 h after IM 1 µg HA saRNA ± 10 µg polySTING ± αIFNAR antibody. *N*=5. Significance calculated via one-way ANOVA test with Tukey’s HSD on raw data. **(b)** Heatmap of Pearson’s correlation matrix of treatment-induced cytokine responses.
